# Synaptic plasticity in the medial preoptic area of male mice encodes social experiences with female and regulates behavior toward young

**DOI:** 10.1101/2023.10.23.560098

**Authors:** Kazuki Ito, Keiichiro Sato, Yousuke Tsuneoka, Takashi Maejima, Hiroyuki Okuno, Yumi Hamasaki, Shunsaku Murakawa, Yuzu Takabayashi, Chihiro Yoshihara, Sayaka Shindo, Haruka Uki, Stefan Herlitze, Masahide Seki, Yutaka Suzuki, Takeshi Sakurai, Kumi O Kuroda, Masabumi Minami, Taiju Amano

## Abstract

A dramatic shift from aggressive infanticidal to paternal behaviors is an essential event for male mice after mating. While the central part of the medial preoptic area (cMPOA) has been shown to critically mediate the paternal behaviors in mice, how this brain region becomes activated by mating and subsequent interaction with pups has not been investigated. Here, we demonstrate that the reduction in inhibitory synaptic strength towards the cMPOA provided by posterior-dorsal medial amygdala (MePD) neurons is a key event for the post-mating behavioral shift in males. Consistent with this, we found optogenetic disinhibition of Me^Cartpt^ to the cMPOA synapses reduces male aggression towards pups. The cMPOA of paternal mice mediated pup-induced neural plastic changes in the bed nucleus of the stria terminalis. These findings provide possible functions of cMPOA neural circuits required for the reception to young in male mice.

## Introduction

Reproductive behavior in male includes courting, mating and parental behavior. Polygynous male mammals, which form reproductive groups with one male and multiple females, often show aggressive behavior toward un-familiar infant conspecifics^1–5^. Infanticide of non-offspring young is commonly believed to improve reproductive success by achieving a shorter inter-birth interval for females, a situation more favorable to breeding lower competition for resources, and the avoidance of misdirected paternal investment^3^,^6^. After mating, however, the same male gradually stops committing infanticide and becomes paternal, even toward non-biological offspring^5^. This behavioral transition from infanticide to paternal care in C57BL/6 male mice is shown to require not only the experience of ejaculation but also cohabitation with the pregnant female^7^,^8^. The exact neural mechanisms governing this behavioral transition through social experiences with female and infant remained unknown.

The medial preoptic area (MPOA) has been identified as one of the most important brain regions regulating parental behavior^9^. Within the MPOA, the central part (cMPOA), located ventral to the cluster of magnocellular oxytocin neurons (the anterior commissural nucleus), appears to play a pivotal role in parental behavior because bilateral cMPOA lesions abolished this behavior and promoted infanticide in both maternal and paternal mice^10^,^11^. Likewise, optogenetic activation of the MPOA in virgin male mice delayed infanticidal behavior^11^,^12^. However, the neural circuit mechanisms controlling the MPOA neuronal activities were unclear. Previously, we reported that behavioral mode transition from infanticide to parental was initiated by lesions of vomeronasal organ (VNO), which transmits olfactory information to the accessory olfactory bulb^8^. Similar results were obtained in the experiment using genetically mutated mice with impaired vomeronasal signaling^12^. Tracing studies using classical retrograde tracer or rabies virus indicate that sensory information from accessary olfactory system is mediated by the bed nucleus of the stria terminalis (BST) and the medial amygdala (Me) and is transmitted to the MPOA^13–15^. It is reported that aggressive behavior toward pup is elicited by activation of GABAergic Me^16^,^17^.

The present study evaluates the efficacy of synaptic transmission from the Me to putative cMPOA neurons from three groups: (i) virgin males, (ii) males that experienced mating and pup exposure, and (iii) an intermediate stage, males that experienced mating but had not pup exposure, referred to as “father in gestation experiences (FGE)” ^18^) (Fig. 1A). We observed that inhibitory inputs from the Me to cMPOA were suppressed before pup delivery, leading to cMPOA disinhibition. Optogenetic inhibition of Me−cMPOA synapses significantly reduced the proportion of mice showing aggressive behavior toward pups. Furthermore, we found paternal experiences increased synaptic inhibition in the rhomboid nucleus of the bed nucleus of the stria terminalis (BSTrh), one of a downstream structure of the MPOA^11^. Together, findings indicate that alterations in plasticity at Me–cMPOA synapses occur as a result of social experience with a female partner, which may function to prime a shift in behavioral response toward pups. This is followed by paternal experience with pups inducing changes in BSTrh.

**Fig. 1.**
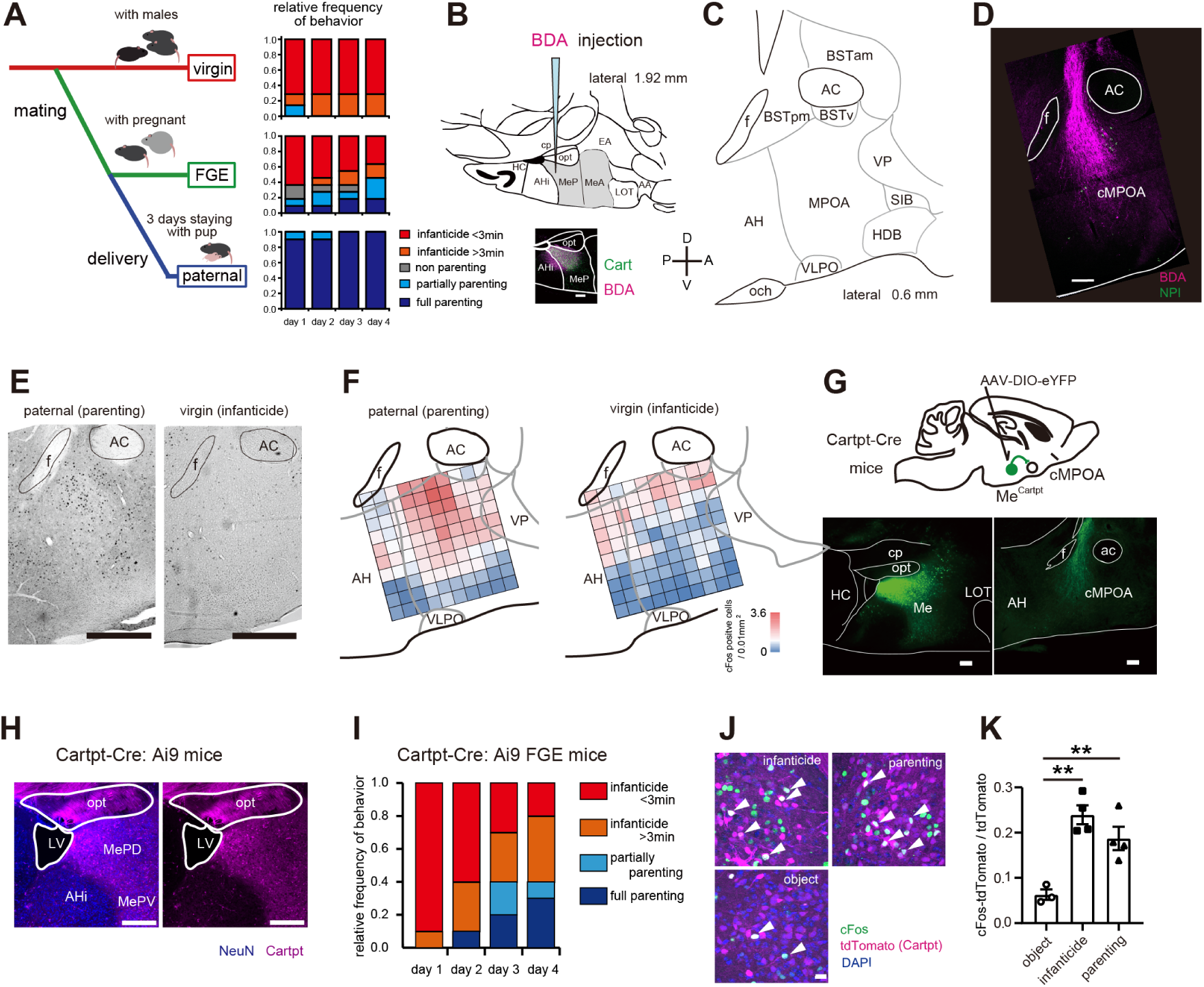
The medial amygdala sends projections to the cMPOA (A) (left) Schematic diagram of “virgin” male mice, intermediate males called “FGE” by mating and cohabitation with partner female, and “paternal” mice, that experienced delivery of partner females and cohabitation with pups for three days. (right) Relative frequencies of behavioral patterns of virgin, FGE, and paternal mice toward pups during four consecutive days of pup exposure. (B) upper, medial amygdala in the parasagittal section referring to the Mouse Brain Atlas of Franklin & Paxinos (2007). lower, Representative images of the Me stained with anti-Cart visualized with Alexa Fluor 488 and BDA visualized with Alexa Fluor 568-conjugated streptavidin (magenta). Scale bar = 200 µm. (C) MPOA in the parasagittal section. (D) Representative images of the Me injected with the tracer BDA in the MPOA. Parasagittal sections are shown. Fixed sections were visualized with Alexa Fluor 568-conjugated streptavidin (magenta). In parallel, fixed sectioned were immunostained with anti-NPI and visualized with Alexa Fluor 647 (green). Scale bar = 200 µm. (E) Image of cFos visualized by DAB staining in the mice showed parenting and infanticide. Scale bar = 500 µm. (F) Expression distribution of cFos-positive neurons in each region delimited by the 100 µm grid. Red-shifted color indicates a higher density of cFos-positive neurons. paternal (n = 8), virgin (n = 3) (G) AAV5-EF1α-DIO-eYFP was injected into the Me of Cartpt-Cre mice. Representative images of eYFP emission from Me and MPOA. Me^Cartpt^ neurons sent projection fibers into the MPOA. Scale bar = 200 µm. (H) Representative images of the MePD in Cartpt-Cre: Ai9 reporter mice. Among MePD neurons immunostained with anti-NeuN antibody visualized with a secondary antibody conjugated to Alexa Fluor 647 (blue), 21.2% Cartpt neurons (magenta) were tdTomato positive (1 male, 1 female). Scale bar = 300 µm. (I) Relative frequencies of behavioral patterns of FGE Cartpt-Cre: Ai9 reporter mice toward pups during four consecutive days of pup exposure. (J) Representative images of cFos induction in the MePD following pup exposure or neutral object to FGE Cartpt-Cre: Ai9 reporter mice. Fixed sections were stained with anti-cFos and visualized with Alexa Fluor 647 (green). Arrowhead indicates the tdTomato-cFos-double-positive neurons. Scale bar = 30 µm. (K) The ratio of c-Fos-positive neurons out of all tdTomato-positive Me^Cartpt^ neurons following pup or object exposure. object (n = 3), infanticide (n = 4), parenting (n = 4). Statistical analysis by one-way ANOVA, **P < 0.01.

## Results

### Cart positive Me neurons project to the cMPOA and respond to pup exposure

To map the projections from the Me to cMPOA, we took advantage of the high expression of the peptide, cocaine and amphetamine regulated transcript (Cart), in the subpopulation of posterior-dorsal Me (MePD) neurons^19^,^20^. Within the Me, clusters of Cart-positive neurons were visualized in the MePD by immunostaining with an anti-Cart antibody (Fig. S1A). We infused biotin dextran amine (BDA), an anterograde neuron tracer, into the Cart-positive area of the Me (Fig. 1B) and observed neural projections descending between fornix and anterior commissure (AC) and entering the cMPOA were observed (Fig. 1C, D). The cMPOA is located ventral to the anterior commissural nucleus, which contains magnocellular oxytocin neurons and thus could be visualized using an antibody for the oxytocin-associated protein neurophysin I (NPI)^10^. In contrast to the Cart-positive area of the Me, the Cart-negative area did not send a dense projection to the cMPOA regardless of tiny BDA leak in the Cart-positive area of the Me (Fig. S1B-E). We observed neural activity marker, cFos expression in MPOA regions receiving projection from Me in Cart-positive regions, with different trends between the paternal and virgin groups, with cFos expression tending to be higher in the paternal group in cMPOA (Fig. 1E, F). Next, we infused an AAV vector encoding Cre-dependent eYFP into the Me of Cartpt (which encodes Cart)-Cre mice for pathway identification, and we observed a clear Cre-dependent eYFP expression in Me^Cartpt^ neurons and eYFP-labeled projections into the cMPOA (Fig. 1G), consistent with the tracer study (Fig. 1B and 1D).

Furthermore, we investigated the neural activity of Me^Cartpt^ neurons after exposure to pups. Cartpt-Cre: Ai9 reporter mice showed that 21.2% of NeuN-positive neurons in MePD were Cartpt (tdTomato) positive (Fig. 1H). We exposed three pups to Cartpt-Cre: Ai9 -FGE mice for four days, and as the days passed, the proportion of mice showing aggressive behavior decreased, while the number of parental individuals increased (Fig. 1I); mice showing aggressive and parental respectively on day four were perfused with PFA 90 min after the end of the behavioral test. The results of the immunohistochemical analysis showed a significant increase of c-Fos positive Cartpt-neurons in the aggressive and parental groups compared to the control group but no difference between the aggressive and parental groups (Fig. 1J, K).

### Sensitivity to GABAergic synaptic inputs from the Me to the cMPOA are suppressed by mating experience and cohabitation with a late gestational female

We infused an AAV vector to express blue light-sensitive channelrhodopsin, ChR2(H134R) in Me^Cartpt^ neurons (Fig. 2A). After incubation periods, we performed whole-cell patch clamp recordings from cMPOA neurons in slices prepared from these mice. Application of blue light to brain slices, including the cMPOA, depolarized the synaptic terminals derived from Me^Cartpt^ neurons and induced both excitatory and inhibitory synaptic currents in cMPOA neurons; as expected, control group mice expressing only eYFP did not show light-evoked synaptic responses (Fig. 2B-D). We evaluated social experience-dependent synaptic changes at both excitatory and inhibitory pathways. At the recording potential (−60 mV), glutamatergic and GABAergic ionotropic receptor currents should flow in the opposite direction. Thus, glutamatergic and GABAergic predominance can be distinguished by the net change in membrane current. Because of the abundance of GABAergic neurons in the MePD^13^, including retrogradely labeled cMPOA projecting MePD neurons (Fig. S2), we chose an experimental protocol that emphasized the EPSC amplitude. The driving force for EPSC and IPSC, calculated by subtraction of the recording potential (−60 mV) and reversal potential (−1 mV for EPSC, −87 mV for IPSC calculated by Nernst equation), was about 59 mV and 27 mV, respectively. Under this experimental protocol, we counted cMPOA neurons expressing predominantly blue light-induced EPSCs or IPSCs by the net membrane current change. Whereas 83.3% of cMPOA neurons were IPSC-dominant in virgin male mice, only 21.4% were IPSC-dominant in FGE males (Fig. 2F). In addition, we observed that intracellular perfusion of a membrane-impermeable G-protein signaling blocker GDPβS into the cMPOA neurons reversed the ratio of IPSC-dominant neurons in FGE males to the level of virgin males. The data presented suggest that experiencing mating and cohousing with a pregnant female changes the excitatory/inhibitory (E-I) balance at synaptic transmission from Me^Cartpt^ into the cMPOA. Whereas application of tetrodotoxin (TTX) blocked light-induced synaptic current, an additional application of 4-aminopyridine rescued, suggesting the monosynaptic projection at the Me^Cartpt^-cMPOA pathway (Fig. 2F).

**Fig. 2.**
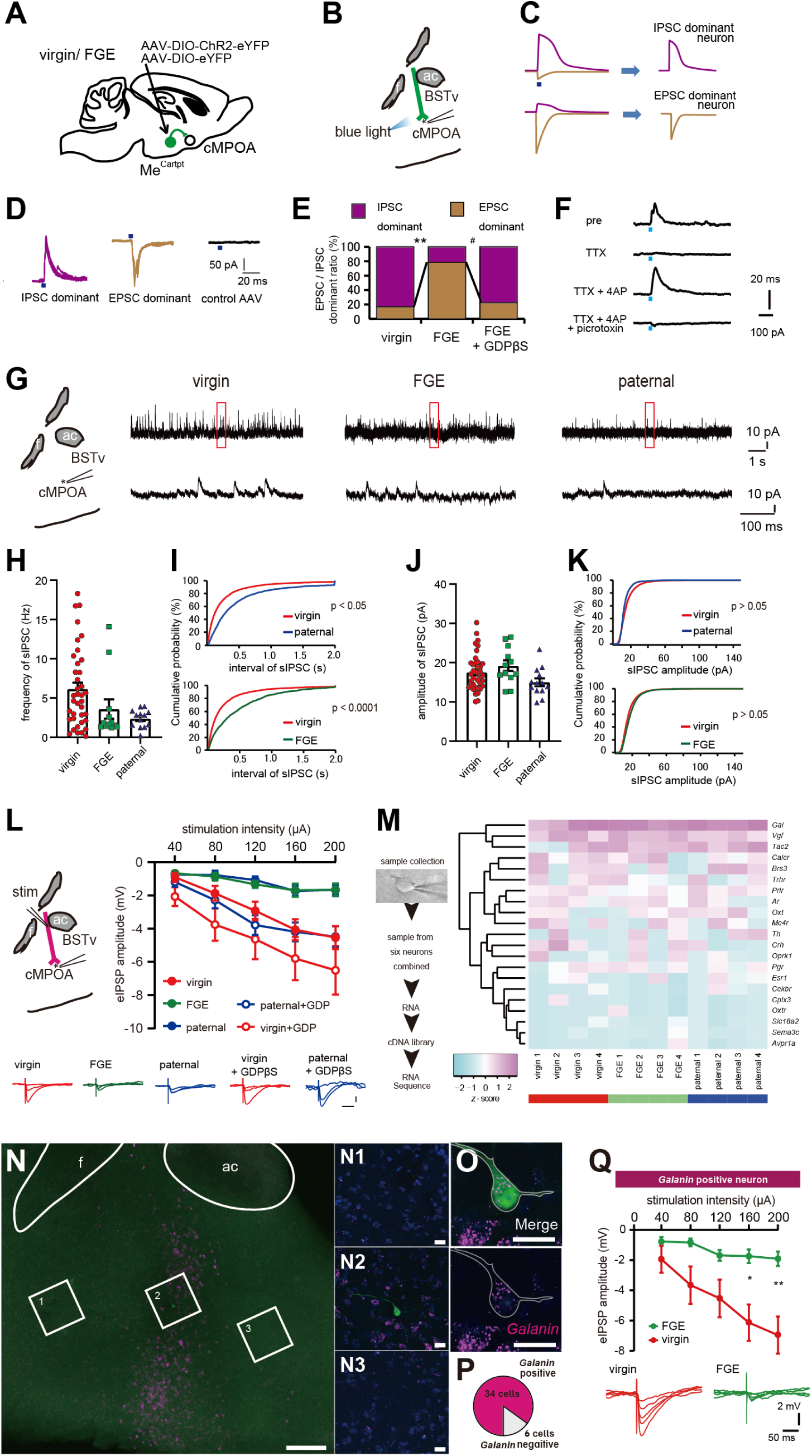
Synapse plastic changes in the cMPOA neuron that occur in the process leading to paternal expression (A) AAV5-EF1α-DIO-eYFP or AAV5-EF1α-DIO-ChR2-eYFP was injected into the Me of Cartpt-Cre mice. After the expression of eYFP with/without ChR2, whole-cell patch clamp recordings were performed from cMPOA neurons in parasagittal sections of virgin and FGE mouse brains. (B) Schematic image of the recording from the cMPOA neuron in a parasagittal section. Input fibers from Me^Cartpt^ expressed ChR2 were stimulated by blue light application. (C) Diagram of dominant responses of IPSC and EPSC in the opposite current direction as that of GABAergic and glutamatergic ionotropic receptors. (D) Representative traces of blue light-evoked IPSCs and EPSCs in IPSC-and EPSC-dominant cMPOA neurons. When Me^Cartpt^ neurons were injected with the control AAV virus, no responses to optical activation were observed in cMPOA neurons. (E) Relative frequencies of excitatory-and inhibitory-dominant cMPOA neurons in virgin mice (n = 12 cells; 4 animals), FGE mice (n = 14 cells; 3 animals), and FGE mice with GDPβS (1 mM) in the pipette solution (n = 9 cells; 2 animals). A significant difference in of excitatory-to inhibitory-dominant cMPOA neurons ratio between virgin and FGE group mice was observed (Fisher’s exact probability test, **P = 0.0048). A significant difference in the ratio of excitatory- and inhibitory-dominant cMPOA neurons between FGE mice and FGE mice with GDPβS (1 mM) in the pipette solution (FGE + GDPβS group) was also observed (Fisher’s exact probability test, #P = 0.0131). (F) Representative traces of blue light-evoked IPSCs in IPSC-dominant cMPOA neurons. The application of TTX blocked light-evoked IPSC. Additional application of 4AP recovered IPSC, which was sensitive to picrotoxin. (G) left, Schematic image of the recording from cMPOA neuron in a parasagittal section. right, Representative traces of sIPSC in cMPOA from virgin, FGE and paternal mice. (H) Frequency of sIPSC in the cMPOA neuron of virgin (n =38 from 11 animals), FGE (n = 12 from 4 animals), and paternal mice (n = 14 from 6 animals). Statistical analysis by Kruskal-Wallis test, *P = 0.0207, (I) Cumulative probability plots of the inter-event interval of the sIPSCs. Statistical analysis by Kolmogorov-Smirnov test, P < 0.0001 (virgin vs FGE), P = 0.0227 (virgin vs paternal), (J) Amplitude of sIPSC in the cMPOA neuron of virgin (n =38 from 11 animals), FGE (n = 12 from 4 animals). Statistical analysis by One way ANOVA, P = 0.0666. (K) Cumulative probability plots of the amplitude of the sIPSCs. Statistical analysis by Kolmogorov-Smirnov test, P > 0.05 (L) left, Schematic image of the recording from the cMPOA neuron in a parasagittal section. Input fibers from MePD were electrically stimulated. right, Input−output curves of stimulus-evoked IPSPs from virgin (n = 18 cells; 6 animals), FGE (n = 14 cells; 4 animals), paternal (n =8 cells; 3 animals), virgin with GDPβS (1 mM) in the pipette solution (n = 13 cells; 4 animals), paternal with GDPβS (n = 16 cells; 4 animals). Statistical analysis by two-way RM ANOVA, P = 0.0016. (M) left, summary of experimental procedure. right, RNA-seq identification of candidate gene expressed in the cMPOA neurons. Heatmap showing a *z*-scored TPM of selected genes for neuronal cluster-mediated social behavior in the MPOA^21^. virgin n = 4 (n = 4), FGE (n = 4), and paternal (n = 4). (N) Representative images of the MPOA. *Galanin* mRNA Alexa Fluor 647 -conjugated hairpin DNA (#S23) (magenta), streptavidin conjugated with FITC (green) and DAPI (blue). (N1-N3) magnified images corresponding area in (N), Scale bar = 200 µm for (N, left) and 20 µm for (N1-N3) (O) Representative images of cMPOA neurons after whole-cell patch-clamp recording. Biocytin was infused from recording pipette. Fixed sections visualized with FITC -conjugated streptavidin (green) and *in situ* HCR for *Galanin* mRNA Alexa Fluor 647 -conjugated hairpin DNA (#S23) (magenta). Scale bar = 20 µm. (P) Ratio of *Galanin* positive cMPOA neurons targeted for whole-cell patch-clamp recording. (Q) Input−output curves of stimulus-evoked IPSPs from the *Galanin* positive cMPOA of virgin mice (n = 12 cells; 7 animals) and FGE mice (n = 7 cells; 5 animals). Values are presented as the mean ± standard error. (bottom) Representative traces used to construct input−output curves (40, 80, 120, 160, and 200 μA stimuli). Statistical analysis by two-way RM ANOVA, P = 0.0397, Holm-Sidak post hoc test, *P < 0.05, **P < 0.01 vs. virgin group.

Such changes in E-I balance could be caused by an increased excitatory or a decreased inhibitory synaptic function. We analyzed spontaneous postsynaptic currents in the cMPOA and found changes in the frequency of spontaneous inhibitory postsynaptic currents (sIPSCs) but no differences in amplitude. A comparison of relative frequencies showed a trend towards less frequent intervals of sIPSC in the paternal and FGE groups than in virgin (Fig. 2H-K). No differences were found in spontaneous excitatory postsynaptic currents (sEPSCs, Fig. S3A-E).

Next, we applied electrical stimulation on the fiber inputs including those from Cart-positive Me to the cMPOA. Similar to FGE mice, paternal mice showed a significant level of decreased eIPSP amplitude in the cMPOA compared with the virgin mice (Fig. 2L). In addition, intracellular infusion of GDPβS reversed eIPSP amplitude in the paternal group. Average evoked excitatory postsynaptic potentials (eEPSPs) amplitudes did not differ between the groups (Fig. S3F). We also addressed the effects of pregnant female exposure alone (i.e., without the experience of copulation and delivery) on virgin males (virgin male with late gestation; VMLG group, Fig. S4A) or sexual experience without staying until late gestation of female partner (male with mating experience; MME, Fig. S4D). As a result, eIPSP amplitude in the cMPOA in VMLG and MME mice was not changed (Fig. S4B and S4E), suggesting that either mating experience or cohabitation with a late gestational female was not enough to reduce eIPSP amplitude in the cMPOA. There were no significant differences in the passive membrane properties of cMPOA neurons among the groups (Table S1). In behavioral test, most mice showed aggression on the first day in both groups; in the VMLG group, this did not change throughout the four days, whereas in the MME group, some mice showed parental behavior as the experimental days passed (Fig. S4C, F).

To address the recorded cell type in the cMPOA, we prepared cell samples using the same experimental methods as patch clamp recordings were manually collected using a glass pipette. RNA-seq analysis was performed to evaluate mRNA expression levels. Six samples were combined and represented as data for one mouse. Table S2 lists the differentially expressed genes (DEGs) in cMPOA neurons among virgin, FGE, and paternal mice. Further, we evaluated the expression levels of 21 representative genes previously reported in activated MPOA neurons during social behavior^21^ to explore the recorded neuron’s cell type in this study and found no difference in their expression levels among the groups. Among all samples, the expression level of *Galanin* (*Gal*) was the highest (Fig. 2M). We visualized *Galanin* mRNA-positive neurons with Alexa Fluor 647 using *in situ* hybridization chain reaction (HCR). We observed a banded *Galanin*-expressing region extending ventrally from the posterior part of the anterior commissure (AC) and covering the cMPOA region (Fig. 2N). Also, the cell types of the recorded neurons in our experimental protocol were addressed by co-staining of post hoc *in situ* hybridization and visualization of biocytin by reacting it with streptavidin conjugated to fluorescein isothiocyanate (FITC). We observed that 85.0% (34 in 40 cells) of the recorded neurons were positive for *Galanin* mRNA co-staining (Fig. 2O, P). When the average evoked inhibitory postsynaptic potential amplitude was compared in the neuron confirmed as *Galanin*-positive, the FGE group showed significantly smaller than the virgin group (Fig. 2Q), as seen in Fig. 2L. These meticulous findings suggest that mating and cohabitation with a female before pup delivery induces synapse plastic changes in the cMPOA of male mice, which densely includes *Galanin*-positive neurons.

### Optogenetic inhibition of Me-to-cMPOA inputs changes the spontaneous synaptic activity

Next, we attempted to silence synapses of Me^Cartpt^ terminals in the cMPOA to examine whether inhibition of these inputs modulates neural and/or synaptic activity. For this, we infused an AAV vector encoding vertebrate long-wavelength opsin (vLWO)^22^,^23^ bilaterally into the Me of virgin Cartpt-Cre mice to suppress Me^Cartpt^ inputs (Fig. 3A). To confirm the inhibitory effects of vLWO, we performed whole-cell patch clamp recordings from Me^Cartpt^ neurons expressing vLWO-eGFP. Application of green light hyperpolarized Me^Cartpt^ neurons (Fig. 3B and 3C) but did not in the control group (Fig. 3D and 3E).

**Fig. 3.**
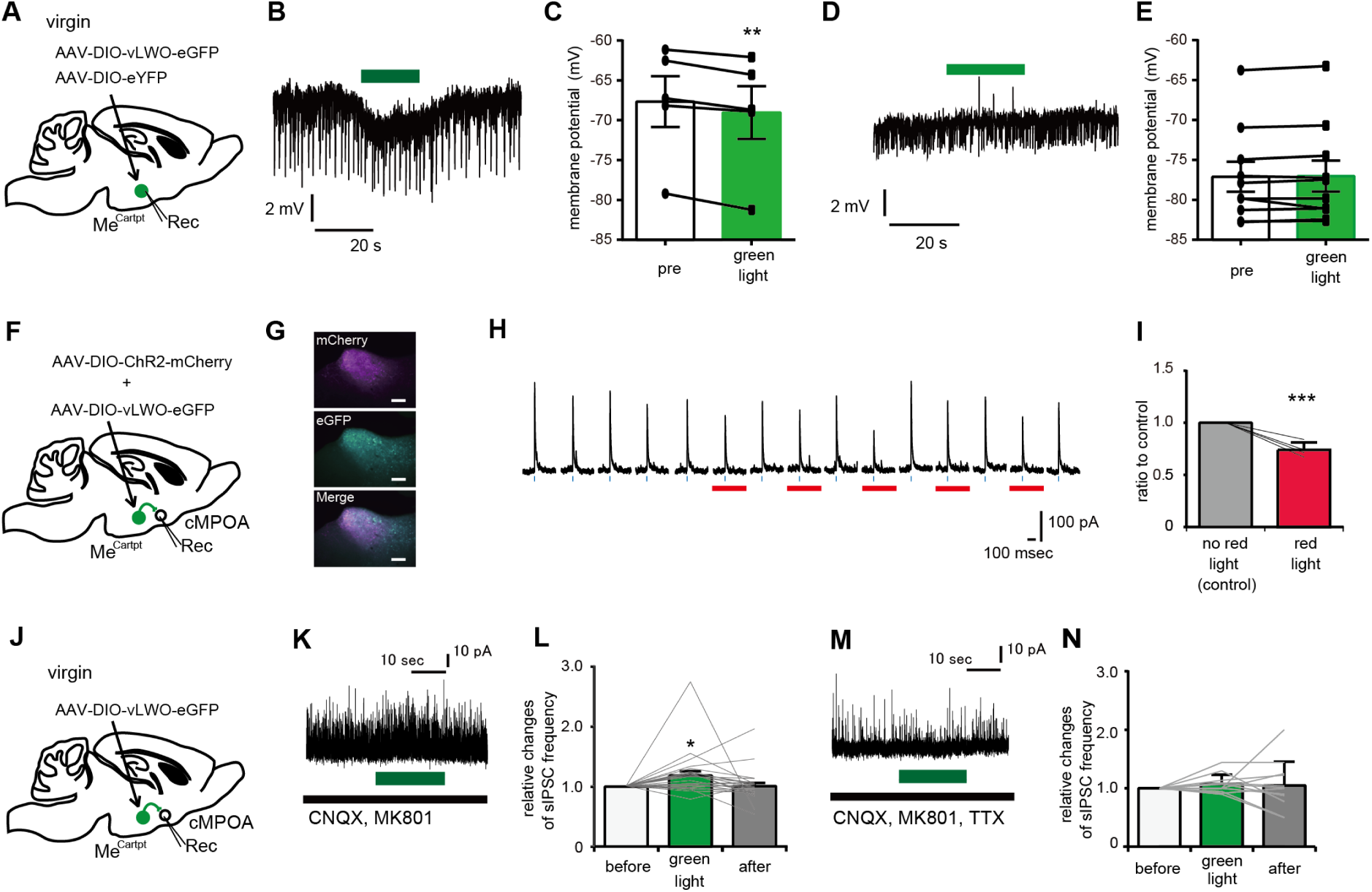
The modulation of Me inputs into the cMPOA (A) AAV10-EF1α-DIO-vLWO-eGFP or AAV10-EF1α-DIO-eYFP were injected into the Me of Cartpt-Cre mice and whole-cell recordings were obtained from fluorescent-labeled Me neurons. (B) Representative resting membrane potential traces recorded from a Me^Cartpt^ neuron expressing vLWO. Green light was applied for 20 s (green bar). (C) Effects of green light application on the resting membrane potential in the Me^Cartpt^ neurons expressing vLWO. The data are shown as the averaged potentials between −6 s and −1 s before the onset of green light application (pre-green light) and between 5 s and 10 s after the onset of green light application (green light). Lines represent the data obtained from individual neurons (n = 5), and bars represent the averaged data obtained from five neurons. Statistical analysis by two-tailed paired t-test, **P = 0.0017, compared with pre-green light application. (D) Representative resting membrane potential traces recorded from a Me^Cartpt^ neuron expressing control eYFP. Green light was applied for 20 s (green bar). Five animals were excluded due to exclusion criteria. (E) Effects of green light application on the resting membrane potential in the Me^Cartpt^ neurons expressing control eYFP. Lines represent the data obtained from individual neurons (n = 10), and bars represent the averaged data obtained from nine neurons. Statistical analysis by two-tailed paired t-test, P = 0.6308, compared with pre-green light application. (F) AAV5-EF1α-DIO-ChR2(H134R)-mCherry and AAV10-EF1α-DIO-vLWO-eGFP-5HT1A was injected into the Me of Cartpt-Cre mice and whole-cell recordings were obtained from cMPOA neurons. (G) Representative images of the Me expressing eGFP and mCherry. Scale bar = 200 µm. (H) Representative traces of blue light-evoked IPSCs in the cMPOA neurons. Red bar indicates red light application (2 s). (I) Effects of red-light application on the blue light-evoked postsynaptic currents in the cMPOA neurons. Lines represent the data obtained from individual neurons (n = 4), and bars represent the averaged data obtained. Statistical analysis by two-tailed paired t-test ***P = 0.0007, compared with pre-red light application. (J) AAV10-EF1α-DIO-vLWO-eGFP was injected into the Me of Cartpt-Cre mice and whole-cell recordings were obtained from cMPOA neurons. (K) Representative trace of green light-induced modulation of sIPSCs in the cMPOA neurons. Green light was applied for 20 s. (L) Effects of green light application on the frequency of sIPSC in the cMPOA neurons. Lines represent the data obtained from individual neurons (n = 25), and bars represent the averaged data. Statistical analysis by repeated measures ANOVA, P = 0.0396, followed by Dunnett’s multiple comparisons post hoc test *P = 0.0332 compared with pre-green light application. One data showed large sIPSC frequency. It was judged as an outlier in Grubbs’ test. Without this data point, the same repeated measure ANOVA analysis resulted in F (1.733, 39.87) = 4.149, P=0.0279, indicating that the conclusion remains when leaving out this outlier. (M) Representative trace of sIPSCs in the cMPOA neurons in the presence of TTX (1 μM). (N) Application of TTX blocked the green light induced changes of IPSCs frequency. Lines represent the data obtained from individual neurons (n = 12), and bars represent the averaged data. Statistical analysis by repeated measures ANOVA, P = 0.6561 followed by Dunnett’s multiple comparisons post hoc test *P = 0.2338 compared with pre-green light application.

Next, we infused an AAV vector encoding ChR2 and vLWO into the Me of virgin Cartpt-Cre mice to confirm the inhibitory effects of vLWO at the synaptic terminal (Fig. 3F and 3G). After more than four weeks, brain slices were prepared and recorded from cMPOA neurons. A green light could affect ChR2, and therefore, instead of using green light as above, the red light was used to stimulate the vLWO. The amplitude of blue light-induced IPSC was significantly reduced by vLWO activation (Fig. 3H and 3I). Furthermore, the suppressed synaptic inputs from the Me^Cartpt^ to the MPOA could result in reduced direct synaptic transmission and in more frequent inhibitory synaptic currents by disinhibition of surrounding interneurons. To address this hypothesis, we observed spontaneous IPSCs (sIPSC) in each cMPOA neuron before and after inhibiting inputs from the Me^Cartpt^ via vLWO activation (Fig. 3J-L). Within 25 recorded neurons, sIPSC frequency was increased by more than 10% in 13 neurons and reduced by more than 10% in 1 neuron. The average frequency of the sIPSC in cMPOA was significantly increased (Fig. 3L). The application of TTX blocked the effects of vLWO activation (Fig. 3M and 3N). These data suggest that inhibition of the Me^Cartpt^ inputs to the cMPOA could modify the activity of each neuron in the cMPOA by acting on an inter-neuronal network.

### In vivo optogenetic inhibition of Me-cMPOA inputs suppresses infanticidal behavior

Next, we aimed to address the causal relationship between synapse plastic changes in the cMPOA and behavioral changes. To address the behavioral causality of the impaired Me-cMPOA inputs, we infused an AAV vector encoding vLWO into the Me of virgin Cartpt-Cre mice, and two optic fibers were bilaterally implanted above the cMPOA for optogenetic stimulation. While the green light was delivered to the cMPOA, three pups were exposed (Fig. 4A). The proportion of vLWO-expressing mice that did not show infanticidal behavior was more significant than in the control group. About this relative frequency of behavioral patterns, we found statistically significant differences between the vLWO and the control group (Fig. 4B). After a four-day test with a green light application, subject mice were exposed to pups again without the green light application. As a result, a similar proportion of subject mice as day four did not infanticide again. The latency to first sniffing did not differ significantly between the vLWO group and control group (Fig. 4C). Latency and probability of each mouse committing infanticide throughout test days were shown in the survival curve (Fig. 4D). Because we detected altered GABAergic neurotransmission in the cMPOA of FGE and paternal mice, we tested the effects of inhibition of the Me input to cMPOA in vGAT-IRES-Cre mice. As a result, similar trends to the case of Carpt-Cre mice were observed in vGAT-IRES-Cre mice with statistically significant differences (Fig. 4E-I). These data suggest that inhibition of the Me inputs, mainly GABAergic, to the cMPOA disrupts aggression of virgin male mice toward pup without minimal effect on investigation motivation.

**Fig. 4.**
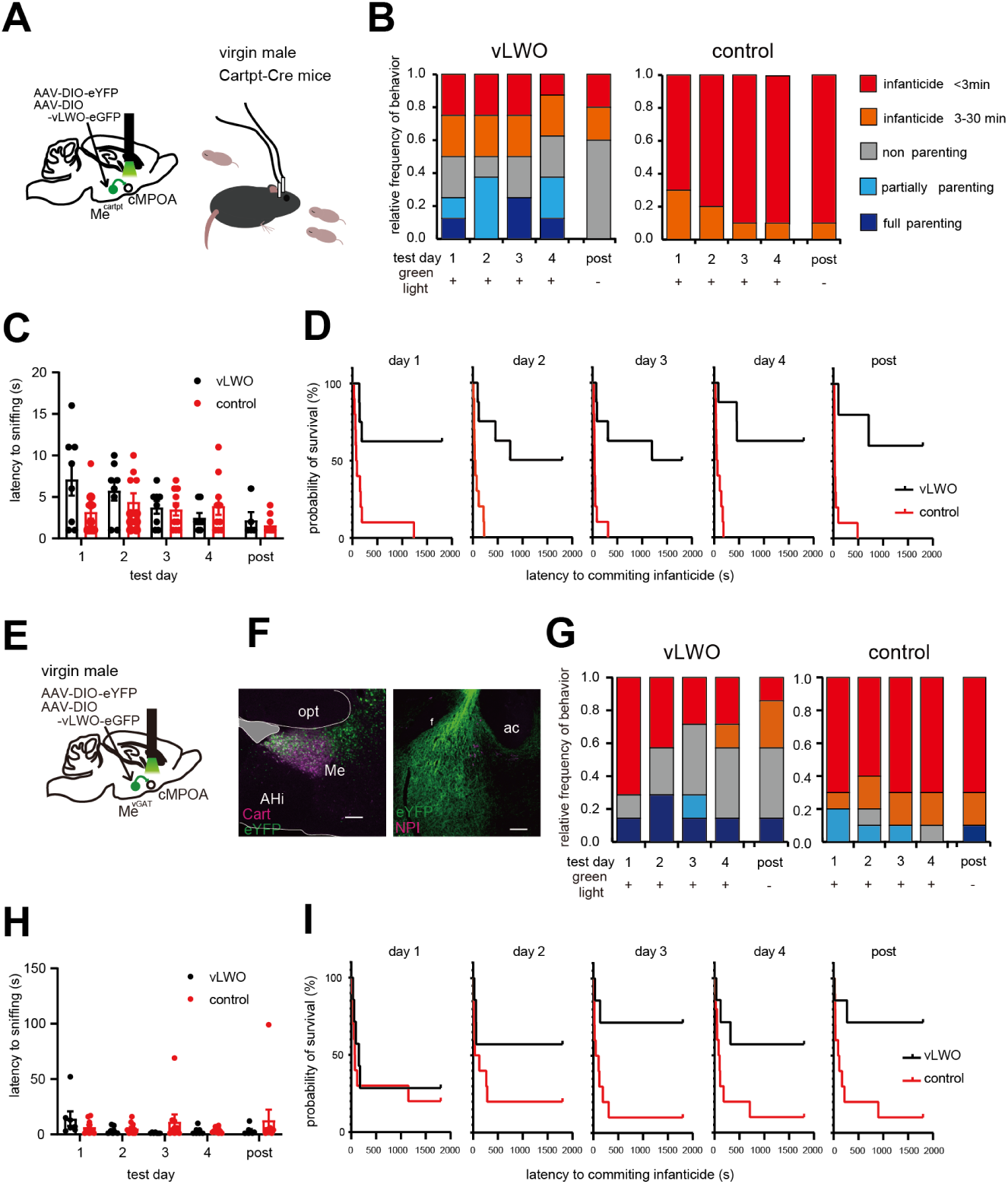
Optogenetic manipulation of the Me^Cartpt^−cMPOA input suppressed infanticidal behavior of virgin male mice (A) AAV10-EF1α-DIO-vLWO-eGFP-5HT1A or AAV10-EF1α-DIO-eYFP was injected into the Me of Cartpt-Cre mice. (B) Relative frequencies of paternal/infanticidal behaviors toward pups during manipulation of the Me^Cartpt^-to-cMPOA input by vLWO activation. Significant difference in the ratio showing infanticidal behavior between mice expressing vLWO-eGFP (n = 8, n =5 for post) or eYFP only (n = 10). Analysis of Variance of Aligned Rank Transformed Data, F (1.236, 13.60) = 63.27, P < 0.0001. (C) Latencies to first sniffing were compared between vLWO and control group mice. Statistical analysis by two-way RM ANOVA, F (1, 16) = 1.642, P = 0.2183. (D) Survival curve to indicate the probability of committing infanticide on each experimental day. vLWO-eGFP (n = 8, n =5 for post) or eYFP only (n = 10). (E) AAV10-EF1α-DIO-vLWO-eGFP-5HT1A or AAV10-EF1α-DIO-eYFP was injected into the Me of vGAT-IRES-Cre mice. (F) Representative images of eYFP expressed in the Me of vGAT-IRES-Cre mice (green). Sections were immunostained with anti-Cart and visualized with a secondary antibody conjugated to Alexa Fluor 594 (magenta). All scale bar = 200 µm. (G) Relative frequencies of paternal/infanticidal behaviors during vLWO activation at the Me^vGAT^ input into cMPOA. Significant difference in the ratio showing infanticidal behavior between mice expressing vLWO-eGFP (n = 8, n =5 for post) or eYFP only (n = 10). Analysis of Variance of Aligned Rank Transformed Data, F (1.236, 13.60) = 5.2140, P = 0.026 (H) Latencies to first sniffing were compared between vLWO and control group mice. Statistical analysis by two-way RM ANOVA, F (1, 16) = 1.642, P = 0.2183. (I) Survival curve to indicate the probability of committing infanticide on each experimental day. vLWO-eGFP (n = 8, n =5 for post) or eYFP only (n = 10).

### Paternal experience enhances inhibitory transmission in the BSTrh through a postsynaptic mechanism

We have demonstrated that the decreased inhibition of MPOA by Me input upon experiences with female is a crucial event for suppressing infanticide in the FGE. However, about half of FGE mice still show infanticidal behavior, whereas almost all paternal mice do not show infanticidal behavior^8^. Next, we examined the synaptic change in paternal mice upon experiences with the pup.

The downstream structure of the MPOA, the rhomboid nucleus of the bed nucleus of the stria terminalis (BSTrh), is activated by infanticidal behavior and negatively regulated by cMPOA^11^. To address the impact of social experiences on the BSTrh synaptic property, we measured the changes in the excitatory—inhibitory (E/I) ratio of the virgin, FGE, and paternal groups in a manner similar to our previous studies^24^. The average amplitude of IPSPs evoked by electrical stimulation of the stria terminalis (Fig. 5A) was enhanced in the BSTrh of the paternal group compared to FGE and virgin groups (Fig. 5B). However, the average eEPSP amplitudes did not differ significantly between these groups (Fig. 5C). These data suggest that, in the paternal mice, increased inhibition in the BSTrh critically occurs following experiences with pups, called pup sensitization.

**Fig. 5.**
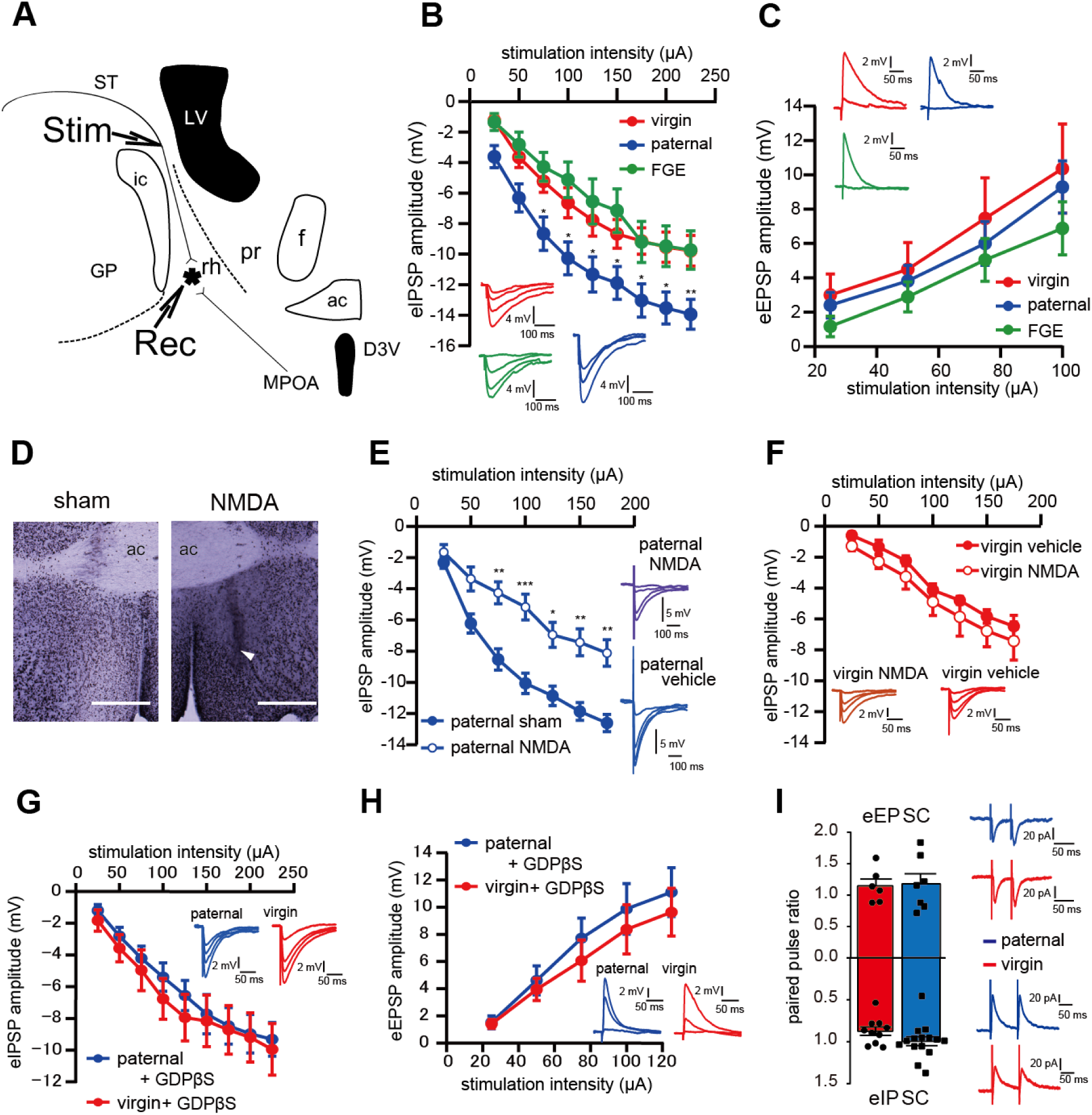
Inhibitory synaptic inputs into the BSTrh were potentiated in paternal group mice (A) Whole-cell patch clamp recordings were performed on BSTrh neurons (Rec). To observe the evoked synaptic responses, electrical stimulation was delivered through a stimulating electrode (Stim). rh, rhomboid nucleus; pr, principal nucleus; LV, lateral ventricle; ic, internal capsule; f, fornix; D3V, dorsal third ventricle; GP, globus pallidus; ac, anterior commissure; ST, stria terminalis (B) Input−output curves of stimulus-evoked IPSPs from virgin mice (n = 16 cells; 10 animals), paternal mice (n = 17 cells; 11 animals), and FGE mice (n = 10 cells; 5 animals). Values are presented as the mean ± standard error. (inset) Representative traces used to construct input−output curves (50, 100, 150, and 200 μA stimuli). Statistical analysis by two-way RM ANOVA, P = 0.0083, followed by Holm-Sidak post hoc test, *P < 0.05, **P < 0.01 vs. virgin group. (C) Input−output curves of stimulus-evoked EPSPs from virgin mice (n = 9 cells; 6 animals), paternal mice (n = 15 cells; 6 animals), and FGE mice (n = 6 cells; 3 animals). Values are presented as the mean ± standard error. (inset) Representative traces used to construct input−output curves (25 and 100 μA stimuli). Statistical analysis by two-way RM ANOVA, P > 0.05. (D) Photographs of the MPOA from vehicle-injected and NMDA-lesioned mice. Sections were immunostained with anti-NeuN and visualized with DAB. Arrowhead indicates the ventral edge of lesioned area. Scale bar = 500 μm. (E) Input−output curves of stimulus-evoked IPSPs from paternal vehicle-injected mice (vehicle; n = 26 from 7 mice, NMDA; n = 15 from 6 mice). Values are presented as the mean ± standard error. (inset) Representative traces used to construct input−output curves (25, 75, 125, and 175 μA stimuli). Statistical analysis by two-way RM ANOVA, P = 0.0032 followed by Holm-Sidak post hoc test, *P < 0.05, **P < 0.01, ***P < 0.001 vs. virgin group. (F) Input−output curves of stimulus-evoked IPSPs from vehicle-injected virgin mice (vehicle; n = 7 cells; 4 animals) and cMPOA-lesioned virgin mice (NMDA; n = 9 cells; 4 animals). Values are presented as the mean ± standard error. (inset) Representative traces used to construct input−output curves (25, 75, 125, and 175 μA stimuli). Statistical analysis by two-way RM ANOVA, P > 0.05. (G) Input−output curves of stimulus-evoked IPSPs from virgin mice (n = 10 cells; 4 animals) and paternal mice (n = 9 cells; 4 animals) recorded with GDPβS (1 mM) in the pipette solution. Values are presented as mean ± standard error. (inset) Representative traces used to construct input−output curves (50, 100, 150, and 200 μA stimuli). Statistical analysis by two-way RM ANOVA, P > 0.05.| (H) Input−output curves of stimulus-evoked EPSPs from virgin mice (n = 9 cells; 5 animals) and paternal mice (n = 10 cells; 4 animals) recorded with GDPβS (1 mM) in the pipette solution. Values are presented as mean ± standard error. (inset) Representative traces used to construct input−output curves (25, 75 and 125 μA stimuli). Statistical analysis by two-way RM ANOVA, P > 0.05. (I) (top) Paired-pulse ratio of eEPSCs (inter-stimulation interval = 50 ms) from virgin mice (n = 6 cells; 5 animals) and paternal mice (n = 7 cells; 6 animals). Statistical analysis by two-tailed unpaired t-test: P > 0.05. (bottom) Paired-pulse ratio of eIPSCs (inter-stimulation interval = 100 ms) from virgin mice (n = 10 cells; 9 animals) and paternal mice (n = 15 cells; 10 animals). Statistical analysis by two-tailed unpaired t-test, P > 0.05.

We hypothesized that the cMPOA plays a critical role in increasing inhibition in the BSTrh. To evaluate this hypothesis, we specifically lesioned the MPOA. To this end, we bilaterally injected NMDA into the cMPOA in virgin and paternal mice (Fig. 5D and S5) and, 3 to 9 days later, performed whole-cell recordings from BSTrh. In the BSTrh of ipsilaterally lesioned paternal mice, the average eIPSP amplitude was significantly smaller than in unlesioned paternal mice (Fig. 5E). These data strongly suggest that the increased inhibitory input to the BSTrh is mediated by the cMPOA in mice showing paternal behavior. It should be noted that cMPOA fiber inputs into BSTrh were not likely stimulated to evoke synaptic responses in the current protocol because cMPOA lesion unchanged eIPSP amplitude in virgin mice (Fig. 5F).

We next addressed whether this increased inhibition onto the BSTrh occurs through a pre-or post-synaptic mechanism. There were no significant differences in passive membrane properties among the three groups (Table S3). We carried out a patch-clamp analysis on BSTrh of paternal group mice. We found out that while inclusion of the GDPβS in the pipette does reduce the amplitude of eIPSPs to the level of the virgin group (Fig. 5G), eEPSP did not show apparent changes (Fig. 5H). These data suggest that the increased eIPSP amplitude in the BSTrh occurs mainly by the postsynaptic mechanisms. We also analyzed the paired-pulse ratios to compare the release probability from presynaptic terminal and found out that the ratio of eEPSCs and eIPSCs did not differ between paternal and virgin groups (Fig. 5I). Altogether, we conclude that the cMPOA in the paternal mice mediates plastic changes at GABA synapses in the BSTrh mainly by the postsynaptic mechanisms, and these sequential events are centrally involved in the fundamental switch in the paternal behaviors of the father animals.

### Effects of pup sensitization on the inhibitory synapse in the cMPOA and in the BSTrh through a postsynaptic mechanism

Repeated exposure of pups to rodents, even male mice, elicits parental behavior^25^,^26^. These results inspired us to investigate synaptic function in male mice that have never mated but engage in parenting behavior. We previously reported that the majority of virgin male C57BL/6J mice before sexual maturation showed retrieval of pups to their nest^24^. Here, virgin male mice at postnatal day 28 experienced pup exposure four times per week and repeated three, six, and eight weeks. No attacks on pups were observed during this period. At 12 weeks old, we tested the behavioral pattern in response to pup exposure (Fig. 6A). As a result, consecutive pup exposures did not result in infanticide on pups beyond sexual maturity; instead, each mouse exhibited pup retrieval to the nest. The proportion of pup-sensitized mice showing infanticide increased in proportion to the length of the gap between the end of pup exposure and the behavioral test (Fig. 6B).

**Fig. 6.**
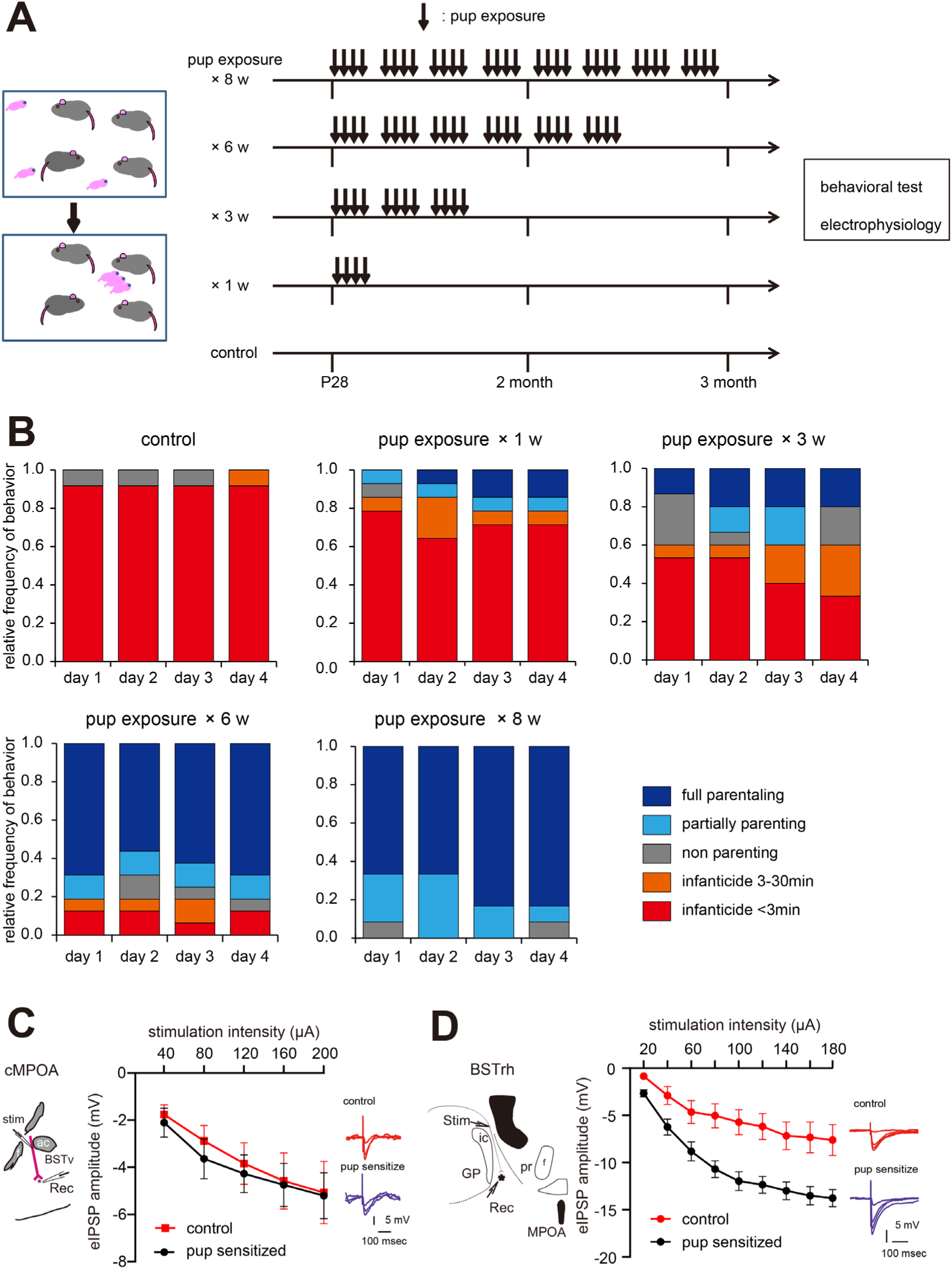
Inhibitory synaptic inputs into the BSTrh were potentiated in pup-sensitized mice (A) Schematic diagram of continuous pup exposure. Four male mice at postnatal day 28 for the first time were exposed to three pups for 30-60 min, four times per week. After one, three, six, and eight weeks of this protocol, mice were used for the experiment. (B) Relative frequencies of paternal/infanticidal behaviors of one, three, six, and eight-week pup exposed and control mice. (C) left, a schematic image of the recording from the cMPOA neuron in a parasagittal section. Input fibers from MePD were electrically stimulated. middle, Input−output curves of stimulus-evoked IPSPs in the cMPOA from pup-sensitized mice (n = 15 cells; 4 animals) and control mice (n = 9 cells; 3 animals). Statistical analysis by two-way RM ANOVA, P = 0.7579, right, representative trace (40, 120, 200 μA stimuli) (D) left, a Schematic image of the recording from the BSTrh neuron in a parasagittal section. Stria terminalis was electrically stimulated. middle, Input−output curves of stimulus-evoked IPSPs in the BSTth from pup-sensitized mice (n = 16 cells; 4 animals) and control mice (n = 9 cells; 5 animals). Statistical analysis by two-way RM ANOVA, P = 0.0016, right, representative trace (20, 60, 100, 140 μA stimuli) Values are presented as the mean ± standard error.

Next, we examined the inhibitory synaptic change in pup-sensitized mice upon experience with the pup for eight weeks. The amplitude of eIPSPs in the cMPOA in pup-sensitized males was similar to that of control males without experience of sensation (Fig. 6C). On the other hand, the amplitude of eIPSPs in the BSTrh in pup ed male were potentiated compared with control males (Fig. 6D) as seen in paternal mice (Fig. 5B). These data suggest that the experiences with pup sensitization encode inhibitory synapse in the BSTrh.

## Discussion

The results showed that multi-step plastic synapse changes correlate with the transition from infanticidal to parental behavior in male mice (Fig. S6). The synapses from Me^Cartpt^ neurons in the cMPOA were altered by interactions with females (Fig. 2). Optogenetic inhibition of the Me^Cartpt^ - cMPOA pathway resulted in impaired aggression of virgin male mice (Fig. 4), suggesting plastic change in Me^Cartpt^ -cMPOA synapses is the potential trigger for the behavioral transition from virgin to FGE. In addition, the shift from FGE to paternal was accompanied by an increased BSTrh inhibition (Fig 5). These plastic changes caused the change of E/I balance, which correlated positively with the neuronal activity of the cMPOA and BSTrh when virgin or paternal males were contacted with pups.

The Me has been implicated in various social behaviors, including social investigation, mating, and aggression^8^,^12,27–30^. In males, the functional association between the Me and the MPOA is critical for mating behavior^31^,^32^. The Me comprises several subdivisions with neural activity patterns, distinct projections and cell types^28^,^33,34^, including the MePD, a sexually dimorphic nucleus crucial for aggressive behaviors^12^,^28,35,36^. In the present study, we focused on Cartpt, which has been reported to be enriched in two of the 16 clusters of the Me^20^. Cartpt-positive neurons were more responsive to pups than to neutral objects (Fig. 1J, K). These data support the importance of investigating the contribution of the Me^Cartpt^-MPOA pathway in the neural mechanisms of social behavior.

Each subpopulation of Me neurons might specifically control some cMPOA neurons and the associated behavioral pattern. Our targeted cMPOA area are mainly composed by *Galanin*-positive neurons (Fig 2M-O), consistent with the previous reports that the activation of *Galanin*-positive MPOA neuron cause inhibition of aggressive behavior toward pup^12^,^14^. The MPOA contributes to several types of intrinsic behaviors, activating the MPOA neurons according to different patterns^11^,^14,28–30,37–39^. However, the mechanisms underlying the expression of one appropriate behavior remain unknown. Suppressing the accessory olfactory signals mediating the Me, presumably changes neural activity patterns in the cMPOA. This hypothesis is supported by the results that temporal disinhibition of the Me—cMPOA pathway modifies behavioral patterns and increases cMPOA inter-neuronal network activity, as reflected by the increased frequency of sIPSCs (Fig. 3). In addition to the Me, other inputs into the cMPOA would drive several inter-neuronal network activity patterns to achieve the multi-functional MPOA and behavioral choices.

In the present study, we show for the first time that social experience with females, including mating, alters the E-I balance in synaptic transmission in the cMPOA in male mice. Changes in GABAergic neurotransmission were observed in sIPSCs and electrical stimulation-induced postsynaptic potentials of FGE and paternal group (Fig. 2L), but not in glutamatergic neurotransmission (Fig. S3). These results were different from the case observed in pregnant female mediating estradiol and progesterone to induce parental behavior^40^. Me^Cartpt^ neurons contain more GABAergic neurons, supporting the possibility of altered GABAergic synaptic function in the cMPOA, the projection site from Me^Cartpt^. In addition, changes in the probability of neurotransmitter release from the presynaptic side and receptor sensitivity and expression levels are known to cause changes in the E-I balance. Me^Cartpt^: Ai9 neurons showed more cFos positive neurons against pups than in neutral objects, but no significant differences between the infanticide and parental groups (Fig. 1K). In the electrophysiological analysis, GDPβS infusion into the recording cMPOA neurons reversed the E-I balance in FGE mice to the level of virgin (Fig. 2C). Thus, it is suggested that changes in GABA_A_ receptor in the cMPOA neurons cause the E-I balance changes in the Me^Cartpt^-cMPOA pathway. However, these hypotheses do not match the reduced sIPSC frequency in FGE mice (Fig. 2G-K); the hypothesis that only some GABA synapses within a single cMPOA neuron are drastically altered may explain the results. It is also necessary to take into account complex mechanisms, such as changes in activity occurring in neurons throughout the MPOA associated with synaptic changes that alter the pattern of intercellular communication, rather than merely input from the Me^Cartpt^. The details require further investigation. How does the experience with female mice bring about these changes? As neither behavior nor synaptic function was altered in the VMLG model (Fig S4), we consider it unlikely that the experience with a pregnant female could direct the brain function and behavior of virgin male mice towards the parental side. Similarly, no inhibitory synaptic changes in the cMPOA were observed in male mice that did not experience mating but came to express parental behavior through pup sensitization (Fig. 6). These data suggest that the synaptic changes in the cMPOA that we detected encode the experience of mating and cohabitation with females but not pup. On the other hand, it was previously reported that the number of males showing infanticide decreased about two weeks after ejaculation without cohabitation with the female partner^7^,^41^. The behavior of males toward pups appears to be controlled by factors triggered by ejaculation, which remain effective for several subsequent weeks, thus preventing infanticide of the male’s own offspring. Several groups have reported that prolactin has the effect of promoting paternal behavior^42^,^43^. However, the possibility that mating-induced prolactin concentration increases may influence subsequent paternal expression has been negatively described in studies using the D_2_ receptor agonist bromocriptine. Post-mating neurotransmission in the brain area related to paternal expression may be altered over time by multiple hormones/neurotransmitters and changes in receptor sensitivity, as has been discussed ^44^. Synaptic functional changes that occur on the way to express fatherhood, such as those found in the present study, may provide clues to elicit new developments in clarifying paternal expression. In this study, the MME model, which experienced mating but separated from female mice, showed parental behavior after several days of pup exposure, although most of them showed infanticide on the first experimental day. Also, the MME model differed from the FGE and paternal groups in the amplitude of eIPSP (Fig S4). The experiences of mating and the social isolation period of 2 weeks after mating might affect synaptic function in the cMPOA and behavioral patterns related to aggression and pup sensitization.

Interestingly, unlike the synaptic changes in the Me^Cartpt^ - cMPOA pathways of FGE mice, an inhibitory shift was observed in the BSTrh E/I balance induced by the caring experience (Fig. 5, 6). What is the biological significance of the increased BSTrh inhibition? Like the previous report^11^, we observed that BSTrh inactivation by hM4Di significantly delayed infanticidal behavior in virgin male mice without parental behavior initiation (Fig. S7). This suggests that BSTrh does not block parental behavior directly. On the other hand, considering the reduced latency for the pup retrieval as the experimental days progressed in FGE and paternal mice (Fig. S8), BSTrh plastic changes could contribute to pup sensitization, which underlies the onset of pup caring. The cMPOA lesions reversed the increased BSTrh inhibition in paternal mice to the level of virgin mice, suggesting that the inhibitory transmission of BSTrh was controlled by cMPOA (Fig. 5D and 55E). The following facts support this: (1) cMPOA neurons project into the BSTrh, and (2) unilateral lesion of the cMPOA increases the number of c-Fos positive neurons in the ipsilateral BSTrh of mice exposed to pups^11^. Because increased BSTrh inhibition in paternal mice is sensitive to intracellular GDPβS infusion (Fig. 5G), as seen in the cMPOA (Fig. 2L), some neurotransmitter or peptide metabotropic receptors may influence BSTrh during and/or after pup sensitization. The cMPOA neurons activated during parental behavior contain many neurotransmitters, including galanin, neurotensin, and tachykinin2^10^,^24^. Moreover, the anterior commissural nucleus, adjacent to the cMPOA, contains the third largest population of magnocellular oxytocin neurons. Therefore, cMPOA neurons or adjacent neurons may directly and persistently control GABA_A_ receptor activity in the BSTrh of paternal mice via metabotropic receptors.

The results of this study show that mating and cohabitation with females and pup sensitization are processed in various parts of the brain, resulting in aggressive male mice going into parenting. Converging data suggest that the cMPOA neurons contribute to infanticide inhibition and parental behavior by modifying the inter-neuronal network and projecting it downstream, including BSTrh. Uncovering the relevant MPOA subgroup for each function requires the analysis of neural circuits. Understanding the modulatory mechanism of synaptic plasticity in the cMPOA and BSTrh may provide evidence for possible interventions in parental stress and subsequent child maltreatment.

## Supporting information

Supplementary Table except for 2 and 6

## Author contributions

TA conceived the study with support from KOK. TA, TS and MM designed the experiments. TA, KI, and KS conducted the behavioral experiments. TA and KI performed stereotaxic surgeries. TA, YT, KI, KS, YH, SM, YT, and SS performed histological experiments. TA and HU performed whole-cell patch clamp recordings. TM, TS and SH developed the AAV virus expressing vLWO. TA and HO prepared sample for RNA sequence and YS, MS and KS analyzed data. TA wrote the manuscript with support from KOK and YT. All authors contributed to editing the manuscript.

## Acknowledgements

This work was supported by Japan Society for the Promotion of Science (JSPS) KAKENHI Grant Number 25713044, 16K19747, 16H06279 (PAGS), 18K07584, 21K07494 (to TA), 26282220 (to KOK), 15K01832 (to TM), 19H03328, 20H05068 (to HO), Suzuken Memorial Foundation (to TA), Narishige Neuroscience Research Foundation (to TA), Nishinomiya Basic Research Fund, Japan (to TA), and the fellowship of Special Postdoctoral Researchers Program at RIKEN (to TA), Hokkaido University, Global Facility Center (GFC), Pharma Science Open Unit (PSOU), funded by MEXT under "Support Program for Implementation of New Equipment Sharing System,” RIKEN Brain Science Institute (2011-2015 to KOK), Deutsche Forschungsgemeinschaft He2471/18 Priority Program (SPP1926), SFB874, SFB1280, DFG2471/21, DFG2471/23 project B10 (to SH). We appreciate Dr. Goichi Miyoshi for critically reading this manuscript prior to submission, Dr. Shigeyoshi Fujisawa and Kazunari Miyamichi for supporting genetically modified mice transportation, RIKEN Research Resource Center for maintenance of animals, Kiyomi Imamura, Terumi Horiuchi, Dr. R. Jude Samulski and the UNC Vector Core for helpful supports.

## STAR Methods

### Animals

The Animal Experiment Committee of the RIKEN Brain Science Institute and Institutional Animal Care and Use Committee at Hokkaido University approved all animal experiments, which were conducted in compliance with the National Institute of Health guidelines for the care and use of laboratory animals.

Animal maintenance and male exposure to different levels of mating and paternal experience Most C57BL/6J male mice were bred at the RIKEN Brain Science Institute and Graduate School of Pharmaceutical Sciences, Hokkaido University unless specially described. For the BDA (Thermo Fisher Scientific) tracer injection into Me (Fig. 1A-C and S1), we utilized C57BL/6J male mice from Japan SLC. Cartpt-Cre mice (Jackson Laboratory, stock number 009615), vGAT-IRES-Cre mice (Jackson Laboratory, stock number 016962) and Ai9 (Jackson Laboratory, stock number 007909) were bred at the Graduate School of Pharmaceutical Sciences, Hokkaido University. We backcrossed vGAT-IRES-Cre mice with C57BL/6J mice at least five generations after arrival from Jackson Laboratory. Mice were housed in individually ventilated cages and provided ad libitum access to water and food and maintained under a 12-h light/dark cycle in cages lined with TEK Fresh Standard bedding (Envigo). Mice were weaned at postnatal day 28 (P28). All mutant heterogenic mice were co-housed with wild-type mice until the experiments. To produce paternal group mice, a virgin male mouse was housed with a female for mating. After delivery, the male stayed with pups for 3 days. All paternal group males experienced one delivery and were used for experiments within 20 days after the birth of pups. If pups did not survive for more than 3 days, the male was not included in the paternal group for experiments. Other virgin male mice were mated and co-housed with a female only until late gestation but not delivery, termed ‘fathers in gestation experience’ (FGE) mice.

### Stereotaxic surgery

BDA (1.25 µg/ dissolved in 12.5 nL 0.1 M phosphate buffer), N-methyl-D-aspartic acid (NMDA, Sigma-Aldrich, 20 mg/ml in saline), and AAV5-EF1α-DIO-hChR2(H134R)-eYFP (6.6 × 10^12^ genome copies/ml, provided by Vector Core at the University of North Carolina (UNC) at Chapel Hill, UNC vector core), AAV5-EF1α-DIO-eYFP (5.6 × 10^12^ genome copies/ml, UNC vector core), AAV10-EF1α-DIO-vLWO-eGFP-5HT1A (3.7× 10^13^ or 1.8 × 10^13^ genome copies/ml, UNC vector core), AAV10-EF1α-DIO-eYFP (1.2 × 10^14^ or 3.7 × 10^13^ genome copies/ml, UNC vector core), rAAV2-EF1a-DO_DIO-tdTomato_EGFP-WPRE (more than 7.0 × 10^12^ vg/ml, Addgene), AAV2-hSyn-HA-IRES-eGFP (3.3 × 10^12^ genome copies/ml, UNC vector core), and AAV2-hSyn-HA-hM4Di(Gi)-IRES-mCitrine (5.6 × 10^12^ genome copies/ml, UNC vector core) (100−400 nL/hemisphere), were administered under anesthesia. Briefly, 2−3-month-old male mice were anesthetized by intraperitoneal (i.p.) sodium pentobarbital (30 mg/kg) or the mixture of medetomidine hydrochloride (0.3mg/kg), midazolam (4mg/kg), butorphanol tartrate (5 mg/kg)^45^, and local subcutaneous (s.c.) lidocaine hydrochloride. The skull was exposed, and holes drilled for stereotaxic injection. Glass capillaries of ∼50 µm tip diameter were filled with oil and backfilled with the test drug or AAV vector. Injection was targeted by referring to the Mouse Brain Atlas of Flanklin & Paxinos (2007) coordinates for the Me (AP −1.5 mm, ML 2.1 mm, DV −5.4 mm and AP −1.2 mm, ML 1.9 mm, DV −5.6 mm), cMPOA (AP 0.1 mm, ML 0.55 mm, DV −5.1 mm), and BSTrh (AP −0.1 mm, ML 1.2 mm, DV −4.2 mm). After injection, the skin was closed with a nylon suture. Mice were kept in single housing 3−4 days for recovery. Mice injected with AAV vectors were then maintained under group housing for more than 4 weeks to allow for expression of vector genes. One week before behavioral testing, optic fibers were implanted into the targeted area as described below.

### Histological analysis

Mice were anesthetized by sodium pentobarbital (50 mg/kg, i.p.) and transcardially perfused with 4% paraformaldehyde (PFA) dissolved in phosphate buffered saline (PBS, pH 7.4). Brains were removed from the skull and post-fixed overnight in PFA at 4 °C, followed by incubation in PBS/20% sucrose for 1 day and PBS/30% sucrose for 1−2 days. Brains were then embedded in O.C.T. Compound (Sakura Finetek) at −80 °C and sectioned at 40 µm using a Cryostat (Leica Biosystems) unless otherwise indicated. Some brains were used to prepare sections for electrophysiological analysis. In the case of BSTrh recording after MPOA lesion, serial slices including MPOA were sectioned at 80 µm in cutting solution (see below) using a Leica VT1200 Semiautomatic Vibrating Blade Microtome (Leica). These slices and the slices including BSTrh used for electrophysiological analysis were fixed in 4% PFA and immunostained.

For fluorescent immunohistochemical staining, brain slices were incubated in PBS containing 0.1% Triton X-100 (PBST) for more than 15 min and then in 0.4% Block Ace (Dainihon-Seiyaku) for an hour. Slices were then incubated at 4 °C overnight in Block Ace solution containing primary antibody. After several rinses in PBST, slices were incubated in PBST containing secondary antibody at room temperature. Table S4 shows antibody list.

To stain the BDA anterograde tracer, sections were incubated with Alexa Fluor 568-conjugated streptavidin (1:500, Thermo Fisher Scientific) for 2 hours. After washout of the secondary antibody or streptavidin, slices were mounted on glass slides with VECTASHIELD Mounting Medium (Vector Laboratories). For 3, 3-diaminobenzidine (DAB) staining, immunolabeled slices were incubated in 0.1 M glycine for 10 min, 0.5% H_2_O_2_ for 30 min, and VECTASTAIN ABC reagent (Vector Laboratories) overnight at 4 °C. After rinsing several times with PBST, the slices were incubated with Vector DAB Substrate (Vector Laboratories) including nickel chloride for about 5 min. The stained slices were mounted on gelatin-coated glass slides and dried. After nissl staining, slides were treated with Softmount (Wako). Labeling of *Galanin* positive neurons followed modified protocol of *in situ* hybridization chain reaction (HCR) using short hairpin DNAs^46^,^47^ using probes listed in Table S5. Fixed brain section re-sliced at 75 µm after patch clamp recording with internal solution containing 1% biocytin (Sigma) and 100 nU RNase inhibitor (Promega), or 30 µm brain section fixed by 4% PFA were soaked in methanol for 10 minutes. After washing with PBST, the sections were prehybridized for 10 min at 37°C in a hybridization buffer containing 10% dextran sulfate, 1× standard saline citrate (SSC), 0.1% Tween 20, 50 μg/ml heparin, 1× Denhardt’s solution. The sections were moved to another hybridization solution containing a mixture of 20 nM split-initiator probes, and incubated overnight at 37°C. For *in situ* HCR, 3 μM hairpin DNA solutions were separately snap-cooled before use. The sections were incubated in amplification buffer (10% dextran sulfate in 5× SSCT) with 60 nM hairpin DNA pairs for 2 hours at 25°C. Then, the samples were washed with PBST three times at room temperature.

Immunofluorescence images were captured using an incident-light fluorescence microscope (Leica DM6000B, Leica), confocal laser-scanning microscope (FV-10, Olympus), or fluorescence microscope (BZ-X700, Keyence). To allow for comparison among trials, the contrast and brightness of all photographs were adjusted linearly and uniformly using Adobe Photoshop CS5 (Adobe Systems) unless specially described. For the lesion study, sections were immunostained with anti-NeuN antibody to confirm cell death in the target area. The loss of NeuN-immunoreactive cells in the NMDA-injected group was compared with the saline group to judge the lesioned area.

### Electrophysiology

Electrophysiological recordings were performed as described previously^24^. The artificial cerebral spinal fluid (ACSF) contained 126 mM NaCl, 2.5 mM KCl, 1.25 mM NaH_2_PO_4_, 1 mM MgCl_2_, 2 mM CaCl_2_, 26 mM NaHCO_3_, and 10 mM glucose (pH 7.3). For a cutting solution, NaCl in ACSF was replaced with the same concentration of choline chloride. Mice were injected with pentobarbital (30 mg/kg, i.p.) followed by transcardiac perfusion of ice-cold cutting solution. The brains were removed, and 230-µm brain slices including the target area prepared in ice-cold choline chloride-based cutting solution using a Leica VT1200 Semiautomatic Vibrating Blade Microtome. Brain slices were stored in warmed ACSF at 32 °C for 20–30 min and then kept at room temperature until recording.

For electrophysiological recordings, slices were superfused with 32 °C–34 °C ACSF at 2–4 mL/min in a chamber mounted on a microscope. Neurons were identified using a 40× or 60× lens and an infrared camera (IR-1000, DEGE-MTI). Glass electrodes (World Precision Instruments) of 4–9 MΩ resistance were used for whole-cell patch clamp recordings in response to electrical stimulation using ∼0.5 MΩ glass electrodes containing ACSF. Recording electrodes were filled with a potassium-based internal solution (132 mM K-gluconate, 3 mM KCl, 10 mM HEPES, 0.5 mM EGTA, 1 mM MgCl_2_, 12 mM Na-phosphocreatine, 4 mM Mg-ATP, 0.5 mM Na-GTP, 0.2-0.3 % biocytin, pH 7.25) and signals were recorded using an Axopatch 700B amplifier (Molecular Devices). Only cells with access resistance <30 MΩ and exhibiting action potentials with >60 mV amplitude evoked by positive current injection were included in the analysis. Whole-cell currents were filtered at 3 kHz. The liquid junction potential (11 mV) was compensated. Stimulating electrodes were placed at the dorsal BSTrh or cMPOA to stimulate stria terminalis. To activate the ChR2-expressing axon terminals, we utilized a LED (465 nm, LEX2-LZ4-B, Brainvision Inc.). To activate the vLWO-expressing neurons, green light generated by a 100 W mercury lamp (U-RFL-T) was applied through a fluorescence band pass filter (520-550 nm) (Olympus). Red light was provided by Fiber-Coupled LED (625 nm, Thorlabs). Evoked synaptic currents and potentials were measured at least three times at the same stimulus intensity and averaged. Responses with action potentials were omitted from the analysis. We recorded evoked excitatory postsynaptic potentials (eEPSPs) and eEPSCs at a holding potential of −85 mV in the presence of 100 µM picrotoxin. eIPSPs were recorded at −65 mV and eIPSCs at −60 mV in the presence of 20 µM CNQX and 20 µM MK-801. In experiments using optogenetic suppression using vLWO, slices were also perfused with 25 µM 9-cis-retinal, 0.1% dimethyl sulfoxide, 0.025% (±) - α-tocopherol and 0.2% bovine serum albumin.

### Behavioral test

The tests of paternal behavior toward pups were performed as described previously^24^. Mice were housed individually in cages containing new purified paper bedding (Alpha-Dri, Shepherd Specialty Papers) and a cotton square (Nestlet, Ancare). After 1–2 days, three pups aged 1–6 days were placed into the cage corner avoiding the nest. Tests were performed once each day for 30 min on 4 successive days during the daytime unless otherwise indicated. The endpoint of the aggressive behavior toward pups to terminate the experiments was 2-s screams or visible wounds on the pup skin. Wounded pups were immediately euthanized. The behavioral score was evaluated according to previous reports as follows: 4 = all pups were retrieved, 3 = 1 or 2 pups were retrieved, 2 = no pup was retrieved, 1 = at least one pup was attacked >3 min after placement in the test cage, and 0 = at least one pup was attacked within 3 min after placement in the test cage.

For c-Fos mapping, the procedure was modified from a previous study^8^. Briefly, mice were exposed to a pup protected inside wire-mesh ball (tea balls, 45 mm diameter; Minex Metal, Tsubame, Japan) with about 10 holes (5 mm diameter) placed in a cage corner avoiding the nest for 30 min. Ninety minutes later, the adult male was anesthetized with pentobarbital (50 mg/kg, i.p.) and transcardially perfused with 4% PFA/PBS for brain isolation as described above. The c-Fos immunoreactive cells were counted automatically by Image J software (National Institutes of Health, Bethesda, MD, USA).

### Optogenetic and chemogenetic manipulation of neural activity in vivo

Zirconia ferrules optical fibers (Thorlabs) of 200 µm diameter core (NA = 0.37) were cut and polished for >70% coefficient of transmission. These constructs were bilaterally implanted into the mouse brain area 1-mm above the target area and fixed with dental cement. After the mouse had recovered from the surgery (one week later), the optic fibers were connected via Ceramic Split Mating with 1×2 Branching Fiber-optic Patch Cords-Glass (0.37 NA, Doric Lenses) to a diode laser (450 nm or 532 nm, Changchun New Industries Optoelectronics Tec.) through a FC/PC connector. Averaged light power at the tips between the bilateral fiber tips was set to 30−40 mW (450 nm) or 10−12 mW (532 nm). Light pulses were controlled by TTL Pulse Generators (Doric Lenses). After connecting the fibers, experimental mice were allowed free actiTon for 15 min. During the last minute of the behavioral test, mice received constant light on for 20 s without light pulses followed by a 10-s off period. This photostimulation protocol was continued throughout the 30-min behavioral experiment. After the behavioral experiments, animals were perfused with 4% PFA to prepare the brain slices. Only animals with bilateral expression of fluorescence labeled fiber and the bilateral optic fibers above targeted area between 0.3-1.0 mm from the targeted area were included for the data analysis. For the chemogentic silencing, we infused CNO (2.0 mg / kg, i.p.) and observed the behavioral pattern 30-35 min later. After the behavioral experiments, we immunostained the brain slices with anti-GFP. Only animals with bilateral expression of fluorescent signal in the targeted area were included for the data analysis.

### RNA-seq analysis

Mouse brain slices were prepared using essentially the same procedures for electrophysiological experiments. Six cMPOA neurons were manually collected by using 1 ∼ 2 MΩ glass electrodes containing ACSF and suspended in cell lysis buffer (0.5% Triton-X100, 0.5 U/µL RNase inhibitor, 5 ng/µL yeast rRNA). The cell samples were immediately frozen and stored at −80°C until starting reverse transcription. Reverse transcription and cDNA library preparation were performed using the Smart-seq2 method^48^ with some modifications. Briefly, 12 µM dT_30_VN primer (5’-AAGCAGTGGTATCAACGCAGAGTACT_30_VN-3) were added into microtubes containing cell lysates (final concentration, 1 µM); then, samples were heat-denatured at 70 °C for 3 min, followed by immediate incubation on ice. Next, samples were mixed with a reverse transcription reaction solution containing locked nucleic acid (LNA)-template switching oligo primer (5’-AAGCAGTGGTATCAACGCAGAGTACrGrG+G-3’, rG; riboguanosines, +G; LNA-modified guanosine, final concentration of 0.6 µM) and reverse transcriptase (SMARTscribe, Clontech, final concentration of 5 U/µL), and incubated at 42°C for 120 min followed by at 70°C for 10 min to terminate the reaction. The resultant cDNA samples were purified using solid phase reversible immobilization (SPRI) beads (AMPure XP, Beckman Coulter). The purified cDNA samples were amplified by PCR using the IS primer (5’-NH_2_-AAGCAGTGGTATCAACGCAGAGT-3’, final concentration of 0.24 µM) and Tks Gflex DNA polymerase (Takara Bio, final concentration of 0.025 U/µL). The amplified cDNA samples were purified by SPRI beads. The quality and quantity of cDNA sample was confirmed by an Agilent 2100 Bioanalyzer using the High sensitivity DNA kit (Agilent Technology, CA, USA).

The cDNA samples were then processed to generate indexed libraries for sequencing using a NEBNext Ultra II DNA Library Prep Kit for Illumina (#E7645, New England Biolabs Inc., Ipswich, MA, USA), according to manufacturer’s instructions. Then, libraries were sequenced for 50-bp single-end reads using HiSeq 3000 (Illumina, San Diego, CA, USA). Output fastq files were aligned to the mouse genome (GRCm38/mm10) using STAR aligner^49^. Library size normalization was performed using R (v.4.1.0) by calculating transcripts per million (TPM). Genes with <5 TPM in all samples were excluded from the analysis. We selected 22 genes for neuronal cluster-mediated social behavior in the MPOA based on previous reports^21^. However, *Oprd1* was excluded because of its low expression. Heatmaps were drawn using gplots (v.3.1.1). Differentially expressed genes (DEGs) between virgin, FGE, and paternal mice were analyzed using DESeq2 (v.1.32.0). Genes with a false discovery rate-adjusted p ≤ 0.05 and a log2 fold change ≥2 in either direction were considered DEGs.

### Statistical analysis

Group means were compared by Fisher’s exact test, the two-tailed t-test, one-way or two-way repeated measures ANOVA followed by a post hoc test as indicated. All statistical tests were two-tailed and conducted with GraphPad Prism software 6 and 9 (GraphPad Software, Inc., La Jolla, CA, USA) or R (v.4.4.0) and ARTool (v.2.2.2). To compare the distribution of behavioral pattern of each mouse during optogenetic manipulation, we performed Aligned Rank Transformation (ART) to handle nonparametric factorial data transformations, and two-way ANOVA analysis. A P-value < 0.05 was deemed statistically significant. Statical data was written in the Fig. legends and Table S6.

### Data availability

All data that support the findings presented in this study are available from the corresponding author upon request. The raw sequences have been deposited in the DNA Data Bank of Japan (DDBJ) under the DDBJ BioProject umbrella with accession number PRJDB8470, and to the DDBJ Read Archive DRA008585 (BioSample ID: SAMD00175991-SAMD00176002).

## References

1. Elwood, R.W. (1977). Changes in Responses of Male and Female Gerbils (Meriones-Unguiculatus) Towards Test Pups during Pregnancy of Female. Anim Behav 25, 46–51. Doi 10.1016/0003-3472(77)90066-5.

2. Hausfater, G., and Hrdy, S.B. (1984). Infanticide: Comparative and Evolutionary Perspectives (Routledge).

3. Hrdy, S.B. (1974). Male-Male Competition and Infanticide among Langurs (Presbytis-Entellus) of Abu, Rajasthan. Folia Primatol 22, 19–58.

4. van Schaik, C.P., and Janson, C.P. (2000). Infanticide by Males and its Implications (Cambridge University Press).

5. vom Saal, F.S., and Howard, L.S. (1982). The regulation of infanticide and parental behavior: implications for reproductive success in male mice. Science 215, 1270–1272.

6. Palombit, R.A. (2015). Infanticide as Sexual Conflict: Coevolution of Male Strategies and Female Counterstrategies. Csh Perspect Biol 7. 10.1101/cshperspect.a017640.

7. vom Saal, F.S. (1985). Time-contingent change in infanticide and parental behavior induced by ejaculation in male mice. Physiol Behav 34, 7–15.

8. Tachikawa, K.S., Yoshihara, Y., and Kuroda, K.O. (2013). Behavioral transition from attack to parenting in male mice: a crucial role of the vomeronasal system. J Neurosci 33, 5120–5126. 10.1523/JNEUROSCI.2364-12.2013.

9. Numan, M. (1974). Medial preoptic area and maternal behavior in the female rat. J Comp Physiol Psychol 87, 746–759.

10. Tsuneoka, Y., Maruyama, T., Yoshida, S., Nishimori, K., Kato, T., Numan, M., and Kuroda, K.O. (2013). Functional, anatomical, and neurochemical differentiation of medial preoptic area subregions in relation to maternal behavior in the mouse. J Comp Neurol 521, 1633–1663. 10.1002/cne.23251.

11. Tsuneoka, Y., Tokita, K., Yoshihara, C., Amano, T., Esposito, G., Huang, A.J., Yu, L.M., Odaka, Y., Shinozuka, K., McHugh, T.J., and Kuroda, K.O. (2015). Distinct preoptic-BST nuclei dissociate paternal and infanticidal behavior in mice. The EMBO journal 34, 2652–2670. 10.15252/embj.201591942.

12. Hong, W., Kim, D.W., and Anderson, D.J. (2014). Antagonistic control of social versus repetitive self-grooming behaviors by separable amygdala neuronal subsets. Cell 158, 1348–1361. 10.1016/j.cell.2014.07.049.

13. Keshavarzi, S., Sullivan, R.K., Ianno, D.J., and Sah, P. (2014). Functional properties and projections of neurons in the medial amygdala. J Neurosci 34, 8699–8715. 10.1523/JNEUROSCI.1176-14.2014.

14. Kohl, J., Babayan, B.M., Rubinstein, N.D., Autry, A.E., Marin-Rodriguez, B., Kapoor, V., Miyamishi, K., Zweifel, L.S., Luo, L., Uchida, N., and Dulac, C. (2018). Functional circuit architecture underlying parental behaviour. Nature 556, 326–331. 10.1038/s41586-018-0027-0.

15. Simerly, R.B., and Swanson, L.W. (1986). The organization of neural inputs to the medial preoptic nucleus of the rat. J Comp Neurol 246, 312–342. 10.1002/cne.902460304.

16. Mei, L., Osakada, T., and Lin, D. (2023). Hypothalamic control of innate social behaviors. Science 382, 399–404. 10.1126/science.adh8489.

17. Chen, P.B., Hu, R.K., Wu, Y.E., Pan, L., Huang, S., Micevych, P.E., and Hong, W. (2019). Sexually Dimorphic Control of Parenting Behavior by the Medial Amygdala. Cell 176, 1206–1221.e1218. 10.1016/j.cell.2019.01.024.

18. Sato, K., Hamasaki, Y., Fukui, K., Ito, K., Miyamichi, K., Minami, M., and Amano, T. (2020). Amygdalohippocampal Area Neurons That Project to the Preoptic Area Mediate Infant-Directed Attack in Male Mice. J Neurosci 40, 3981–3994. 10.1523/JNEUROSCI.0438-19.2020.

19. Broberger, C. (1999). Hypothalamic cocaine-and amphetamine-regulated transcript (CART) neurons: histochemical relationship to thyrotropin-releasing hormone, melanin-concentrating hormone, orexin/hypocretin and neuropeptide Y. Brain Res 848, 101–113.

20. Wu, Y.E., Pan, L., Zuo, Y., Li, X., and Hong, W. (2017). Detecting Activated Cell Populations Using Single-Cell RNA-Seq. Neuron 96, 313–329 e316. 10.1016/j.neuron.2017.09.026.

21. Moffitt, J.R., Bambah-Mukku, D., Eichhorn, S.W., Vaughn, E., Shekhar, K., Perez, J.D., Rubinstein, N.D., Hao, J., Regev, A., Dulac, C., and Zhuang, X. (2018). Molecular, spatial, and functional single-cell profiling of the hypothalamic preoptic region. Science 362. 10.1126/science.aau5324.

22. Masseck, O.A., Spoida, K., Dalkara, D., Maejima, T., Rubelowski, J.M., Wallhorn, L., Deneris, E.S., and Herlitze, S. (2014). Vertebrate cone opsins enable sustained and highly sensitive rapid control of Gi/o signaling in anxiety circuitry. Neuron 81, 1263–1273. 10.1016/j.neuron.2014.01.041.

23. Soya, S., Takahashi, T.M., McHugh, T.J., Maejima, T., Herlitze, S., Abe, M., Sakimura, K., and Sakurai, T. (2017). Orexin modulates behavioral fear expression through the locus coeruleus. Nat Commun 8, 1606. 10.1038/s41467-017-01782-z.

24. Amano, T., Shindo, S., Yoshihara, C., Tsuneoka, Y., Uki, H., Minami, M., and Kuroda, K.O. (2017). Development-dependent behavioral change toward pups and synaptic transmission in the rhomboid nucleus of the bed nucleus of the stria terminalis. Behav Brain Res 325, 131–137. 10.1016/j.bbr.2016.10.029.

25. Cai, W.Q., Ma, H., Xun, Y.F., Hou, W.J., Wang, L.M., Zhang, X.N., Ran, Y.F., Yuan, W., Guo, Q.Q., Zhang, J., et al. (2021). Involvement of the dopamine system in paternal behavior induced by repeated pup exposure in virgin male ICR mice. Behavioural Brain Research 415. ARTN 113519 10.1016/j.bbr.2021.113519.

26. Rosenblatt, J.S. (1967). Nonhormonal Basis of Maternal Behavior in Rat. Science 156, 1512-+. DOI 10.1126/science.156.3781.1512.

27. Kondo, Y. (1992). Lesions of the medial amygdala produce severe impairment of copulatory behavior in sexually inexperienced male rats. Physiol Behav 51, 939–943.

28. Allen, W.E., DeNardo, L.A., Chen, M.Z., Liu, C.D., Loh, K.M., Fenno, L.E., Ramakrishnan, C., Deisseroth, K., and Luo, L. (2017). Thirst-associated preoptic neurons encode an aversive motivational drive. Science 357, 1149–1155. 10.1126/science.aan6747.

29. Sano, K., Nakata, M., Musatov, S., Morishita, M., Sakamoto, T., Tsukahara, S., and Ogawa, S. (2016). Pubertal activation of estrogen receptor alpha in the medial amygdala is essential for the full expression of male social behavior in mice. Proc Natl Acad Sci U S A 113, 7632–7637. 10.1073/pnas.1524907113.

30. Liu, H.X., Lopatina, O., Higashida, C., Fujimoto, H., Akther, S., Inzhutova, A., Liang, M., Zhong, J., Tsuji, T., Yoshihara, T., et al. (2013). Displays of paternal mouse pup retrieval following communicative interaction with maternal mates. Nat Commun 4, 1346. 10.1038/ncomms2336.

31. Dominguez, J., Riolo, J.V., Xu, Z., and Hull, E.M. (2001). Regulation by the medial amygdala of copulation and medial preoptic dopamine release. J Neurosci 21, 349–355.

32. Asanuma, M., Nishibayashi, S., Kondo, Y., Iwata, E., Tsuda, M., and Ogawa, N. (1995). Effects of single cyclosporin A pretreatment on pentylenetetrazol-induced convulsion and on TRE-binding activity in the rat brain. Brain Res Mol Brain Res 33, 29–36.

33. Choi, G.B., Dong, H.W., Murphy, A.J., Valenzuela, D.M., Yancopoulos, G.D., Swanson, L.W., and Anderson, D.J. (2005). Lhx6 delineates a pathway mediating innate reproductive behaviors from the amygdala to the hypothalamus. Neuron 46, 647–660. 10.1016/j.neuron.2005.04.011.

34. Pardo-Bellver, C., Cadiz-Moretti, B., Novejarque, A., Martinez-Garcia, F., and Lanuza, E. (2012). Differential efferent projections of the anterior, posteroventral, and posterodorsal subdivisions of the medial amygdala in mice. Front Neuroanat 6, 33. 10.3389/fnana.2012.00033.

35. Morris, J.A., Jordan, C.L., and Breedlove, S.M. (2008). Sexual dimorphism in neuronal number of the posterodorsal medial amygdala is independent of circulating androgens and regional volume in adult rats. J Comp Neurol 506, 851–859. 10.1002/cne.21536.

36. Unger, E.K., Burke, K.J., Jr., Yang, C.F., Bender, K.J., Fuller, P.M., and Shah, N.M. (2015). Medial amygdalar aromatase neurons regulate aggression in both sexes. Cell Rep 10, 453–462. 10.1016/j.celrep.2014.12.040.

37. Chung, S., Weber, F., Zhong, P., Tan, C.L., Nguyen, T.N., Beier, K.T., Hormann, N., Chang, W.C., Zhang, Z., Do, J.P., et al. (2017). Identification of preoptic sleep neurons using retrograde labelling and gene profiling. Nature 545, 477–481. 10.1038/nature22350.

38. McHenry, J.A., Otis, J.M., Rossi, M.A., Robinson, J.E., Kosyk, O., Miller, N.W., McElligott, Z.A., Budygin, E.A., Rubinow, D.R., and Stuber, G.D. (2017). Hormonal gain control of a medial preoptic area social reward circuit. Nat Neurosci 20, 449–458. 10.1038/nn.4487.

39. Murakami, G. (2016). Distinct Effects of Estrogen on Mouse Maternal Behavior: The Contribution of Estrogen Synthesis in the Brain. PLoS One 11, e0150728. 10.1371/journal.pone.0150728.

40. Ammari, R., Monaca, F., Cao, M.R., Nassar, E., Wai, P., Del Grosso, N.A., Lee, M., Borak, N., Schneider-Luftman, D., and Kohl, J. (2023). Hormone-mediated neural remodeling orchestrates parenting onset during pregnancy. Science 382, 76–81. 10.1126/science.adi0576.

41. Elwood, R.W. (1985). Inhibition of Infanticide and Onset of Paternal Care in Male-Mice (Mus-Musculus). J Comp Psychol 99, 457–467. Doi 10.1037//0735-7036.99.4.457.

42. Smiley, K.O., Brown, R.S.E., and Grattan, D.R. (2022). Prolactin Action Is Necessary for Parental Behavior in Male Mice. J Neurosci 42, 8308–8327. 10.1523/JNEUROSCI.0558-22.2022.

43. Stagkourakis, S., Smiley, K.O., Williams, P., Kakadellis, S., Ziegler, K., Bakker, J., Brown, R.S.E., Harkany, T., Grattan, D.R., and Broberger, C. (2020). A Neuro-hormonal Circuit for Paternal Behavior Controlled by a Hypothalamic Network Oscillation. Cell 182, 960-+. 10.1016/j.cell.2020.07.007.

44. Freeman, M.E., Kanyicska, B., Lerant, A., and Nagy, G. (2000). Prolactin: structure, function, and regulation of secretion. Physiol Rev 80, 1523–1631. 10.1152/physrev.2000.80.4.1523.

45. Kawai, S., Takagi, Y., Kaneko, S., and Kurosawa, T. (2011). Effect of three types of mixed anesthetic agents alternate to ketamine in mice. Exp Anim 60, 481–487.

46. Tsuneoka, Y., Atsumi, Y., Makanae, A., Yashiro, M., and Funato, H. (2022). Fluorescence quenching by high-power LEDs for highly sensitive fluorescence in situ hybridization. Front Mol Neurosci 15, 976349. 10.3389/fnmol.2022.976349.

47. Tsuneoka, Y., and Funato, H. (2020). Modified in situ Hybridization Chain Reaction Using Short Hairpin DNAs. Front Mol Neurosci 13, 75. 10.3389/fnmol.2020.00075.

48. Picelli, S., Faridani, O.R., Bjorklund, A.K., Winberg, G., Sagasser, S., and Sandberg, R. (2014). Full-length RNA-seq from single cells using Smart-seq2. Nat Protoc 9, 171–181. 10.1038/nprot.2014.006.

49. Dobin, A., Davis, C.A., Schlesinger, F., Drenkow, J., Zaleski, C., Jha, S., Batut, P., Chaisson, M., and Gingeras, T.R. (2013). STAR: ultrafast universal RNA-seq aligner. Bioinformatics 29, 15–21. 10.1093/bioinformatics/bts635.

