## Supplementary Table except for 2 and 6 for "Synaptic plasticity in the medial preoptic area of male mice encodes social experiences with female and regulates behavior toward young"

Table S1

Passive membrane properties of cMPOA neurons from virgin, FGE, and paternal mice. Values are presented as the mean ± standard error. No significant differences were found for all parameters (One-way ANOVA, P > 0.05).

|  | virgin | | | FGE | | | paternal | | |
| --- | --- | --- | --- | --- | --- | --- | --- | --- | --- |
| resting membrane potential (mV) | -71.96 | ± | 0.67 | -75.00 | ± | 1.82 | -72.92 | ± | 1.10 |
| input resistance (MΩ) | 349.28 | ± | 20.56 | 385.05 | ± | 44.46 | 408.50 | ± | 38.57 |
| threshold (mV) | -65.23 | ± | 0.55 | -65.11 | ± | 1.25 | -63.10 | ± | 0.88 |
| time constant (ms) | 28.23 | ± | 1.84 | 32.62 | ± | 3.55 | 34.36 | ± | 3.33 |
| n | 46 | | | 13 | | | 25 | | |

Table S2

Please refer the Excel file.

Table S3

Passive membrane properties of BSTrh neurons from virgin, FGE, and paternal mice. Values are presented as the mean ± standard error. No significant differences were found for all parameters (One-way ANOVA, P > 0.05).

|  | virgin | | | FGE | | | paternal | | |
| --- | --- | --- | --- | --- | --- | --- | --- | --- | --- |
| resting membrane potential (mV) | -75.64 | ± | 1.10 | -78.44 | ± | 1.22 | -75.19 | ± | 0.58 |
| input resistance (MΩ) | 498.54 | ± | 31.48 | 509.53 | ± | 37.98 | 506.81 | ± | 32.82 |
| threshold (mV) | -62.12 | ± | 0.77 | -61.50 | ± | 0.88 | -60.34 | ± | 0.58 |
| time constant (ms) | 36.97 | ± | 3.08 | 44.20 | ± | 4.40 | 43.40 | ± | 3.17 |
| n | 25 | | | 16 | | | 32 | | |

Table S4

Antibody list

| primary antibody | |  | secondary antibody |  |
| --- | --- | --- | --- | --- |
| NeuN | Millipore, mouse, monoclonal, MAB377 | 1:1000 - 1:10000 | anti mouse IgG biotin conjugated | 1:2000 |
|  |  |  | anti-mouse IgG Alexa 594 and 647 | 1:1000 |
| c-Fos | Calbiochen, polyclonal, PC38 | 1:8000 | anti rabbit IgG biotin conjugated | 1:2000 |
|  | Synaptic systems, rabbit, monoclonal, code # Rb108B5 | 1:1000 | anti-rabbit IgG Alexa 647 | 1:1000 |
| NPI | Santa Cruz, goat, polyclonal, sc-7810 | 1:6000 | anti-goat IgG Alexa 647 | 1:300 |
|  |  |  | anti-goat IgG Alexa 488 | 1:1000 |
| Cart | Phoenix Pharmaceuticals, rabbit, polyclonal, H-003-62 | 1:10000 | anti-rabbit IgG Alexa 488 | 1:1000 |
|  |  |  | anti-rabbit IgG Alexa 594 | 1:1000 |
| GFP | Medical & Biological Laboratories, rabbit, polyclonal, code # 598 | 1:10000 | anti rabbit IgG biotin conjugated | 1:2000 |
| ERα | Millipore, rabbit, polyclonal, code # 06-935 | 1:5000 | anti-rabbit IgG Alexa 488 | 1:1000 |

Table S5

Probe list

Hairpin sequence and conjugated dye

| Hairpin ID | | Sequence | Fluorophore |
| --- | --- | --- | --- |
| S23 | H1 | ATACGACTTCGACGACCACC-CAACTTGAATGGGTGGTCGTCG | Alexa 647 |
|  | H2 | GGGTGGTCGTCGAAGTCGTA-TCGACGACCACCCATTCAAGTT |  |

Split-initiator probe sequences

| Gal-1P1S23 | GGGTGGTCGaaAGCCTAGCAGGATAACGCTGCCTCT |
| --- | --- |
| Gal-2P1S23 | GGGTGGTCGaaGGCCCAGAAGGTAGCCAGCGCTGTT |
| Gal-3P1S23 | GGGTGGTCGaaCGGACAATGTTGCTCTCAGGCAGGG |
| Gal-4P1S23 | GGGTGGTCGaaTTCTCTAGGTCTTCTGAGGAGGTGG |
| Gal-5P1S23 | GGGTGGTCGaaCAGACGATTGGCTTGAGGAGTTGGC |
| Gal-1P2S23 | CTGACAGGGTCACAACCAACAGGAGaaTCGAAGTCGTAT |
| Gal-2P2S23 | ATGATCTGTGGTTGTCAATGGCATGaaTCGAAGTCGTAT |
| Gal-3P2S23 | AAGAAACTGAGAAACTCCATTATAGaaTCGAAGTCGTAT |
| Gal-4P2S23 | CGTGCACAGTGGACATGGTCTCAGGaaTCGAAGTCGTAT |
| Gal-5P2S23 | ATCATAACACAGCTTCAAAGCAGAGaaTCGAAGTCGTAT |

Table S6

Please refer the Excel file.

Abbreviations

| AA | anterior amygdaloid area |
| --- | --- |
| AC | anterior commissure. |
| AH | anterior hypothalamic area |
| AHi | amygdalohippocampal area |
| BST | the bed nucleus of the stria terminalis |
| BSTam | the bed nucleus of the stria terminalis, medial division, anteromedial part |
| BSTpm | the bed nucleus of the stria terminalis, medial division, posterior part |
| BSTrh | the bed nucleus of the stria terminalis, rhomboid part |
| BSTv | bed nucleus of the stria terminalis, ventral part |
| cMPOA | central part of medial preoptic area |
| cp | cerebral peduncle |
| D3V | dorsal third ventricle |
| EA | extended amygdala |
| f | fornix |
| HC | hippocampus |
| HDB | nucleus of the horizontal limb of the diagonal band |
| ic | internal capsule |
| LOT | nucleus of the lateral olfactory tract |
| LV | lateral ventricle |
| MeA | medial amygdaloid nucleus, anterior part |
| MeP | medial amygdaloid nucleus, posterior part |
| MePD | posterior-dorsal medial amygdala |
| MePV | posterior-ventral medial amygdala |
| MPOA | medial preoptic area |
| och | optic chiasm |
| opt | optic tract |
| SIB | substantia innominata, basal part |
| VLPO | ventrolateral preoptic nucleus |
| VP | ventral pallidum |
