## Supplementary Figure S1-8 for "Synaptic plasticity in the medial preoptic area of male mice encodes social experiences with female and regulates behavior toward young"

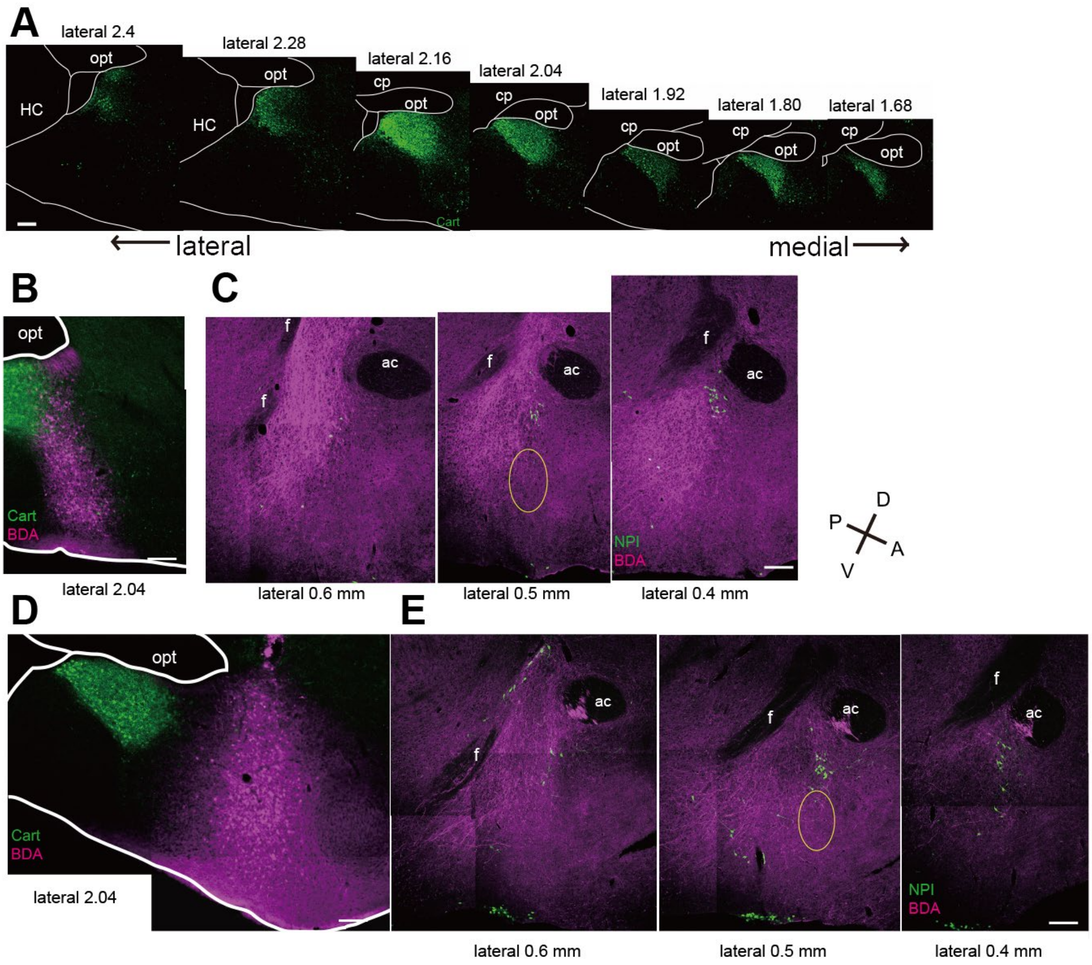


Figure S1

Projection from Cart-negative area in the Me to cMPOA was not dense.

(A) Images of immunostaining with anti-Cart antibody visualized with a secondary antibody conjugated to Alexa 488.

(B) Representative images of the Cart-negative posterior Me injected with BDA. Fixed sections were visualized with Alexa 568-conjugated streptavidin (magenta) and immunostained with anti-Cart visualized with a secondary antibody conjugated to Alexa 488 (left).

(C) The projection to MPOA originating in the posterior Cart-negative Me. Fixed sections were immunostained with anti-NPI (right) and visualized with Alexa 488 (green). Main targeted area as the cMPOA was represented as the area of yellow line.

(D) Representative images of the anterior Me injected with BDA. Fixed sections were visualized with Alexa 568-conjugated streptavidin (magenta) and immunostained with anti-Cart visualized with a secondary antibody conjugated to Alexa 488 (left).

(E) The projection to the MPOA originating from the anterior Me. Fixed sections were immunostained with anti-NPI (right) and visualized with Alexa 488 (green). All scale bar = 200 μm. Main targeted area as the cMPOA was represented as the area of yellow line.

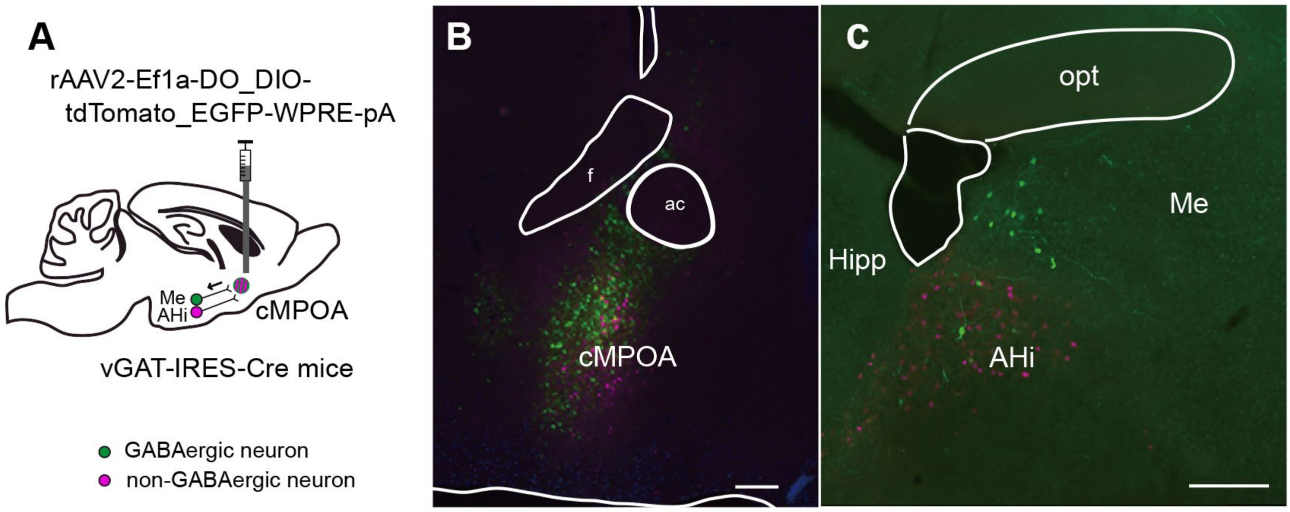


Figure S2

MePD neurons projecting to the cMPOA were mostly GABAergic.

(A) Retrograde virus vector rAAV2-Ef1a-DO_DIO-tdTomato_EGFP-WPRE were injected into the cMPOA of vGAT-IRES-Cre mice. The cMPOA projecting vGAT-positive and vGAT-negative neurons were labeled with EGFP and tdTomato, respectively.

(B, C) Representative images of vGAT positive neurons (green) and vGAT negative neurons (magenta) in the MPOA (virus injection site, B) and Me and surrounding area (C). Whereas putative GABAergic neurons in the MePD were retrogradely labeled with EGFP, more non-GABAergic neuron in the amygdalohippocampal area (AHi) were retrogradely labeled with tdTomato, as reported previously[1]. n = 9, Scale bar = 200 μm.

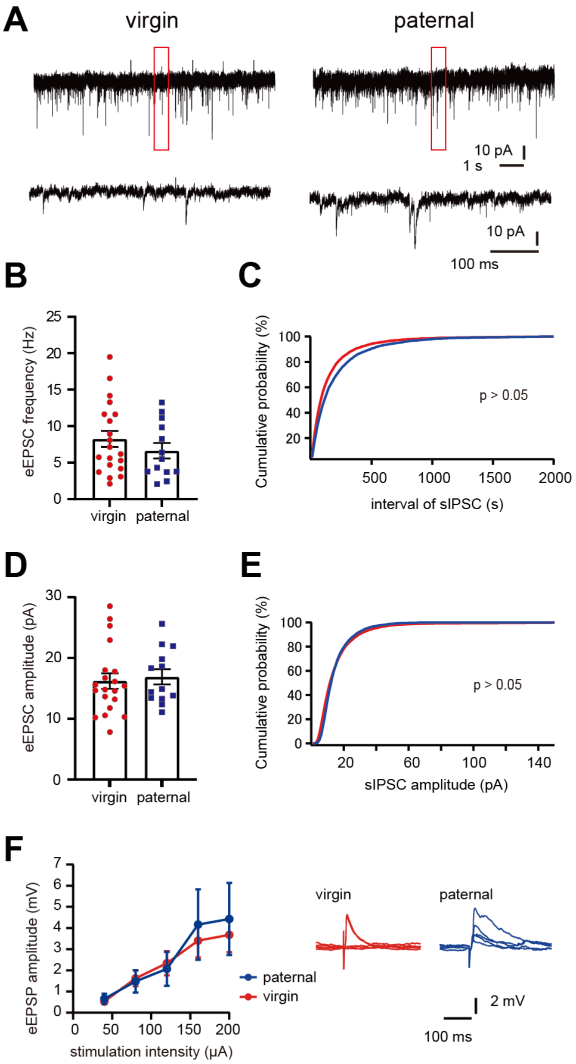


Fig S3

(A) Representative traces of sEPSC in cMPOA from virgin, FGE and paternal mice.

(B) Frequency of sEPSC in the cMPOA neuron of irgin (n =21 from 5 animals), and paternal mice (n = 13 from 4 animals). Statistical analysis by two-tailed unpaired t-test, P = 0.3236

(C) Cumulative probability plots of the inter-event interval of the sEPSCs. Not significant, Kolmogorov-Smirnov test.

(D) Amplitude of sEPSC in the cMPOA neuron of virgin (n =21 from 5 animals), and paternal mice (n = 13 from 4 animals). Statistical analysis by two-tailed unpaired t-test, P = 0.7135

(E) Cumulative probability plots of the amplitude of the sEPSCs. Not significant, Kolmogorov-Smirnov test.

(F) Input–output curves of stimulus-evoked eEPSPs from virgin (n = 21 cells; 3 animals) and paternal (n = 14 cells; 3 animals). Statistical analysis by two-way RM ANOVA, P = 0.7974.

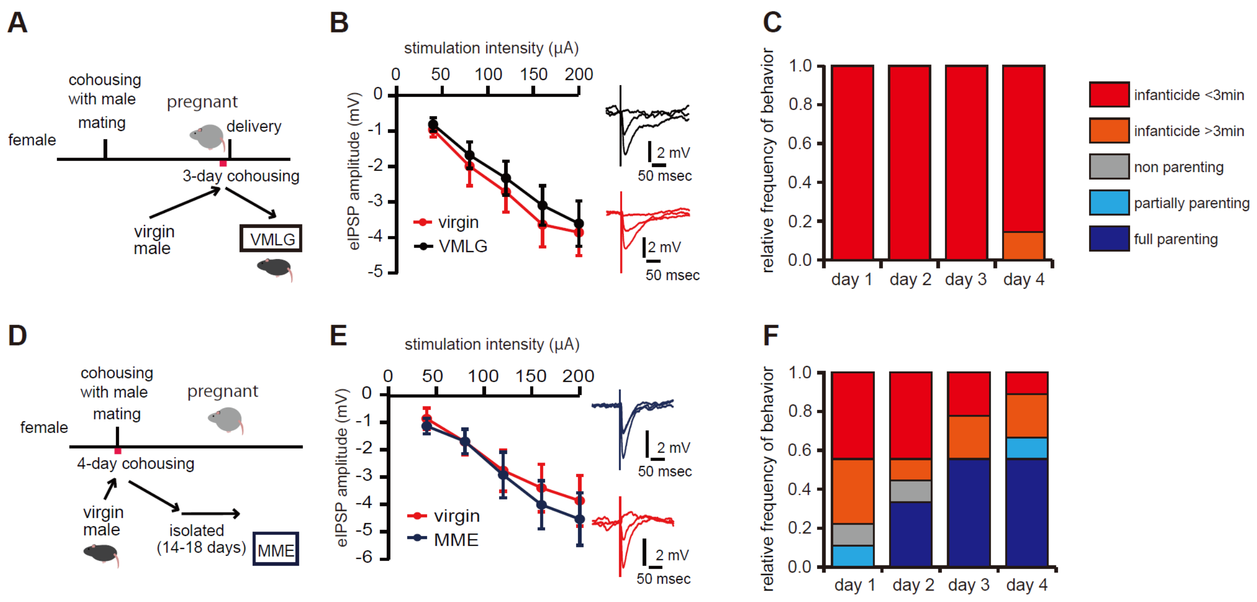


Figure S4

The effects of social experiences with female on eIPSP amplitude in the cMPOA

(A) Establishments of the VMLG mice. Virgin male mice were co-housed for 3 days with a female in late gestation.

(B) Co-housing with pregnant female did not result in the significant plastic changes of cMPOA inputs. Input–output curves of stimulus-evoked IPSPs in the cMPOA from VMLG mice (n = 22 from 4 animal) and virgin mice (n = 19 from 4 animal). Statistical analysis by two-way RM ANOVA, P = 0.5994.

(C) Relative frequencies of behavioral patterns of VMLG mice toward pups (n = 7).

(D) Establishments of the MME mice. Virgin male mice were co-housed for 4 days with female mice and isolated. After 14-18 days, the male mice whose partner female got pregnant were used for the electrophysiological experiments.

(E) Mating experiences without co-housing with pregnant female did not result in the significant plastic changes of Me-to-cMPOA inputs. Input–output curves of stimulus-evoked IPSPs in the cMPOA from MME mice (n = 9 from 4 animal) and virgin mice (n = 11 from 4 animal). Statistical analysis by two-way RM ANOVA, P = 0.7145

(F) Relative frequencies of behavioral patterns of MME mice toward pups (n = 9).

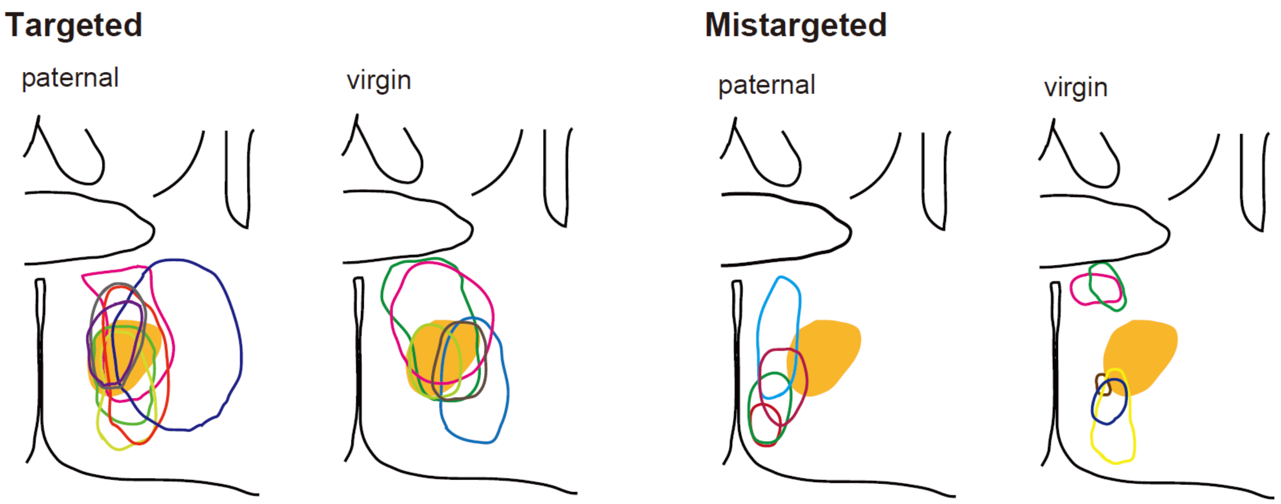


Figure S5

Areas showing damage following NMDA infusion into the MPOA of paternal and virgin mice. Schematic representation of the targeted to the cMPOA (orange), with reference to previous reports[2] (n = 7 for paternal, n = 4 for virgin) and mistargeted to the cMPOA (n = 4 for paternal, n = 5 for virgin) is shown. Images of coronal brain sections that were 0.02 mm anterior to bregma according to Franklin and Paxinos (2007).

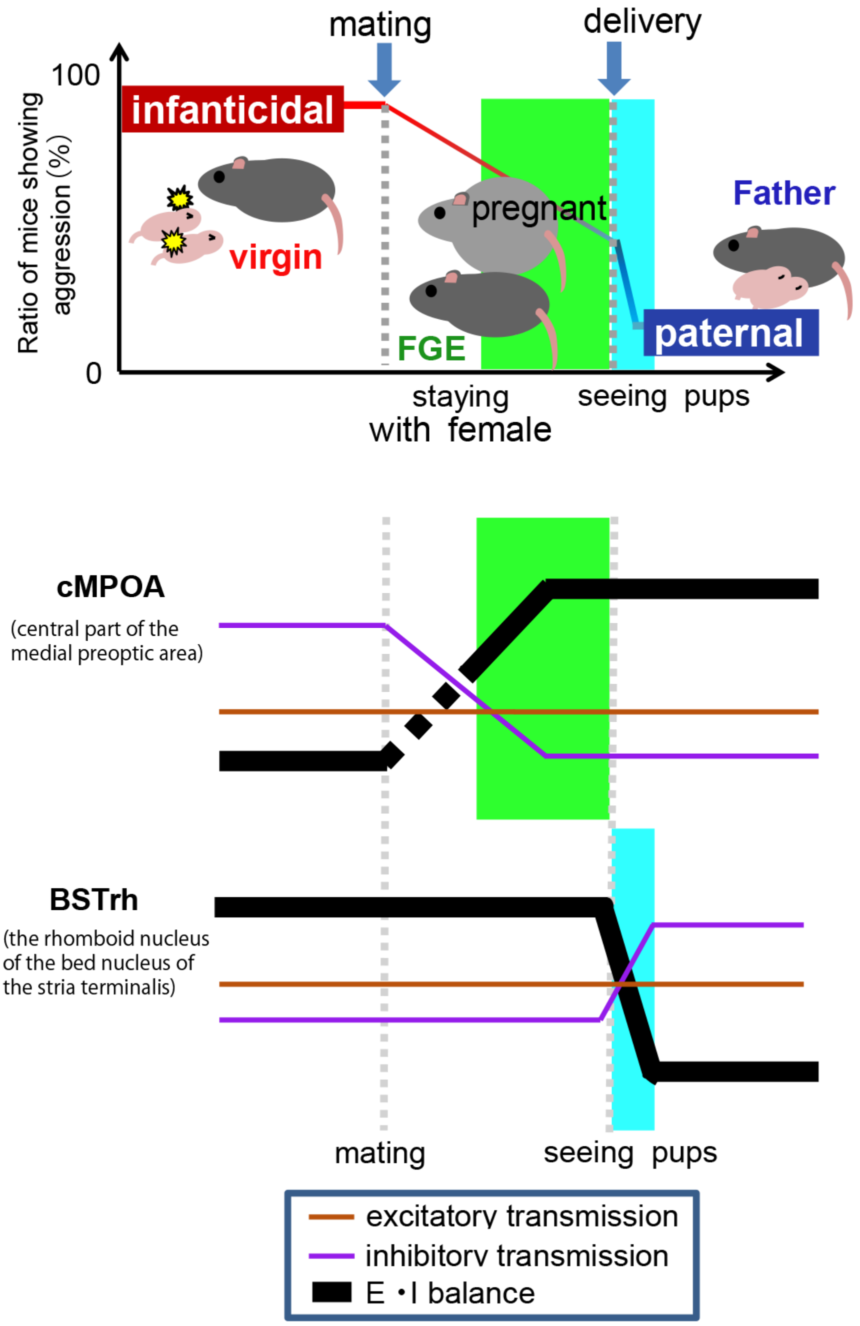


Figure S6

Diagram of the correlation between the synaptic level in the cMPOA and BSTrh and behavioral pattern toward pups

Virgin male mice show infanticide. The ratio of infanticidal male mice was gradually decreased after mating experience. Most of the paternal mice did not show infanticide after experiencing the delivery of pups (top). The E/I balance was shifted toward excitatory in the cMPOA preceding the shift toward inhibitory in the BSTrh. Synaptic input from the Me to the cMPOA was impaired by experience with a female partner (middle, light green). In contrast, synaptic inputs into the BSTrh were potentiated by the experiences of parenting pups (bottom, light blue).

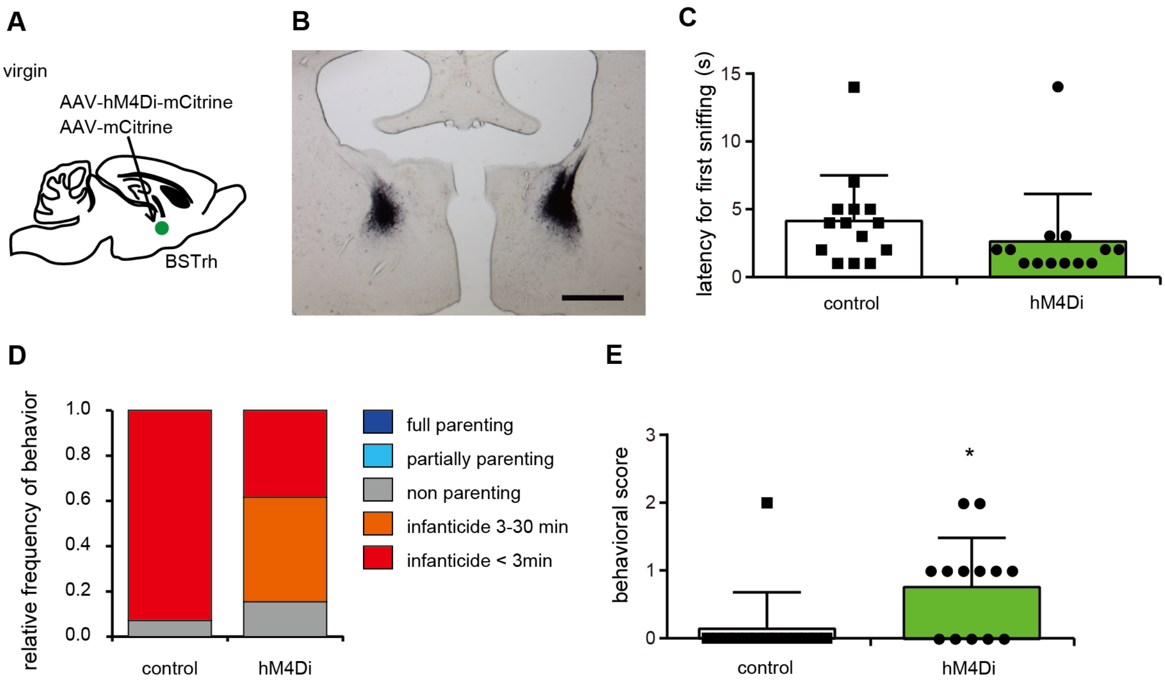


Figure S7

Transient inhibition of BSTrh neurons impaired the infanticidal behavior.

(A) AAV2- EF1α-DIO-hM4Di-mCitrine or AAV2- EF1α-DIO- eGFP (control) was injected into the BSTrh of virgin C57Bl/6J male mice and incubated for four to five weeks. We observed the behavioral pattern 30-35 min after clozapine-N-oxide (CNO; 2.0 mg / kg, i.p.) were injected.

(B) Representative image of injected site. Sections were immunostained with anti-GFP and visualized with secondary antibody conjugated with biotin and DAB. Scale bar = 500 μm.

(C) Latencies to first sniffing were compared between hM4Di (n= 13) and control (n = 14) groups were compared for one day. Statistical analysis by unpaired t-test compared with control group, P = 0.2591

(D) Relative frequencies of paternal or infanticidal behaviors toward pups. Significant difference in the ratio showing infanticidal behavior within 3 min and others between mice expressing hM4Di (n = 13) or control (n = 14). Statistical analysis by Fisher's exact probability test, **P = 0.0044. Eighteen animals of hM4Di group were excluded due to exclusion criteria.

(E) Behavioral scores of hM4Di (n= 13) and control (n = 14) groups were compared. Statistical analysis by Mann-Whitney test **P = 0.01.

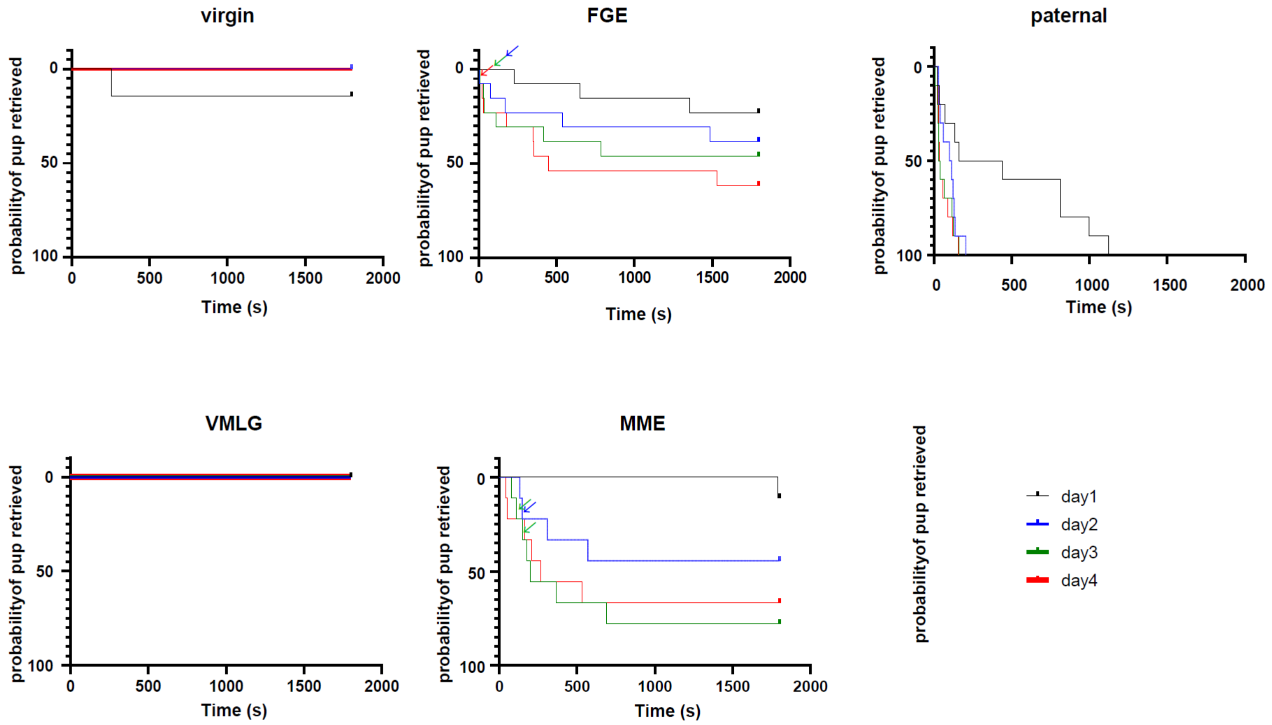


Figure S8
Probability of first pup retrieving of virgin (n = 7), FGE (n = 13), paternal (n = 10), VMLG (n = 7), and MME (n = 9) mice on four consecutive days. Arrowhead indicate the mice showed infanticidal behavior following retrieving.

1 Sato, K. *et al.* Amygdalohippocampal Area Neurons That Project to the Preoptic Area Mediate Infant-Directed Attack in Male Mice. *J Neurosci* **40**, 3981-3994 (2020). https://doi.org/10.1523/JNEUROSCI.0438-19.2020

2 Tsuneoka, Y. *et al.* Distinct preoptic-BST nuclei dissociate paternal and infanticidal behavior in mice. *The EMBO journal* **34**, 2652-2670 (2015). https://doi.org/10.15252/embj.201591942
